# Genomic Architecture of Cells in Tissues (GeACT): Study of Human Mid-gestation Fetus

**DOI:** 10.1101/2020.04.12.038000

**Authors:** Feng Tian, Fan Zhou, Xiang Li, Wenping Ma, Honggui Wu, Ming Yang, Alec R. Chapman, David F. Lee, Longzhi Tan, Dong Xing, Guangjun Yin, Ayjan Semayel, Jing Wang, Jia Wang, Wenjie Sun, Runsheng He, Siwei Zhang, Zhijie Cao, Lin Wei, Shen Lu, Dechang Yang, Yunuo Mao, Yuan Gao, Kexuan Chen, Yu Zhang, Xixi Liu, Jun Yong, Liying Yan, Yanyi Huang, Jie Qiao, Fuchou Tang, Ge Gao, X. Sunney Xie

## Abstract

By circumventing cellular heterogeneity, single cell omics have now been widely utilized for cell typing in human tissues, culminating with the undertaking of human cell atlas aimed at characterizing all human cell types. However, more important are the probing of gene regulatory networks, underlying chromatin architecture and critical transcription factors for each cell type. Here we report the Genomic Architecture of Cells in Tissues (GeACT), a comprehensive genomic data base that collectively address the above needs with the goal of understanding the functional genome in action. GeACT was made possible by our novel single-cell RNA-seq (MALBAC-DT) and ATAC-seq (METATAC) methods of high detectability and precision. We exemplified GeACT by first studying representative organs in human mid-gestation fetus. In particular, correlated gene modules (CGMs) are observed and found to be cell-type-dependent. We linked gene expression profiles to the underlying chromatin states, and found the key transcription factors for representative CGMs.

**Highlights:**

- Genomic Architecture of Cells in Tissues (GeACT) data for human mid-gestation fetus
- Determining correlated gene modules (CGMs) in different cell types by MALBAC-DT
- Measuring chromatin open regions in single cells with high detectability by METATAC
- Integrating transcriptomics and chromatin accessibility to reveal key TFs for a CGM

## Introduction

A human individual cell, as the basic biological unit of our bodies, carry out its functions through rigorous regulation of gene expression, exhibit heterogeneity among each other in every human tissue. Single-cell sequencing technologies have allowed us to characterize genomic profiles (e.g. genome, transcriptome, methylome, chromatin architectures and 3D structures) of individual cells, and have become the most effective way of cell typing, i.e. categorizing each cell type by its genomic features.

Single-cell RNA-seq by next-generation sequencers, since its inception (Tang et al., 2009), has been rapidly advanced by high-throughput development (Klein et al., 2015; Macosko et al., 2015) and widely applied to overcome the cellular heterogeneity, which is particularly suited for tissue samples contain multiple cell types (Cao et al., 2019a; Pijuan-Sala et al., 2019; Wen and Tang, 2019). This prompted the emergence of cell atlases of different organisms, including humans by virtue of cell typing (Cao et al., 2019a; Han et al., 2018; Tabula Muris Consortium, 2018).

Although current scRNA-seq methods have led to discoveries of new and rare cell types, their low RNA detectability limited the number of detected genes in each individual cell. In general, existing methods reporting only expression levels of genes provided little information about gene-gene interactions and regulatory networks. Such information would be available through pairwise correlations between any two genes, but remains unmeasurable with the low RNA detectability (Chapman et al., 2020).

Recently MALBAC-DT (see Methods) has improved RNA detectability, allowing not only more genes to be detected, but also the covariance matrix of all expressed genes, yielding the correlated gene modules (CGMs), i.e. clusters of intercorrelated genes that carry out certain biological functions together. It was found in cell lines that genes within a CGM have a higher probability for protein-protein interactions (Chapman et al., 2020). However, whether CGMs exist in human tissues remains uncharted.

ATAC-seq was first developed to identify genome-wide accessible chromatin regions (Buenrostro et al., 2013), which are critical for the regulation of gene expression. Chromatin accessible regions are cell-type-specific (Cusanovich et al., 2015). Single-cell ATAC-seq has been widely used for cell typing, creating the *cis*-regulatory maps of the whole organism such as *Drosophila* and mouse (Cusanovich et al., 2018a; Cusanovich et al., 2018b). However, scATAC-seq has been conducted with limited detectability, resulting in false negatives of accessible regions in a single cell (Buenrostro et al., 2018; Cusanovich et al., 2018a; Preissl et al., 2018). It is highly desirable to have such a map for humans with low dropout rate.

In this work, we used a novel high-detectability method named METATAC (Xie et al., 2018) (see Methods), which exhibited a ∼100-fold increase in unique DNA fragments from accessible chromatin regions compared with the previous method (Cusanovich et al., 2018a). We used it to generate a chromatin accessibility map of different human tissues, together with the MALBAC-DT data.

Chromatin open regions are accessible by transcription factors (TFs) (Buenrostro et al., 2013), which regulate gene expression, program cell functions, dictate cell differentiation and development (Lambert et al., 2018). Although binding motifs for TFs are available from the database derived from ChIP-seq data, most of them are false positive binding sites according to the futility theory (Wasserman and Sandelin, 2004). Having the CGM and genome architecture at the same time, we could delineate the key TFs associated with the CGMs.

Powered with the two newly developed single-cell techniques (MALBAC-DT and METATAC), we set out to determine GeACT for human tissues. Here, as the first application, we present human mid-gestation (19-21-week) fetuses (Table S1), during which the human fetus undergoes massive organ development and maturation. To the best of our knowledge, this has not been reported previously.

We profiled well-curated transcriptomic and chromatin accessibility landscapes of multiple organs across the digestive, immune, circulatory, respiratory, reproductive, and urinary systems. We identified hundreds of CGMs, co-expressed in one or more cell types. Integrative analyses in two modalities offer a unique opportunity to find cell-type-specific *cis*-regulatory elements for a particular gene, to quantify contributions of the open-chromatin architecture to gene expression, and furthermore, to identify key transcription factors responsible for each CGM.

All the gene expression/activity data, computational tools and pipelines in this study are publicly released on the website at http://geact.gao-lab.org. The precise mapping drafted here will be of important reference values for the study of diseases related to human organ development, carrying potential clinical applications.

## Results

### Construction of the single-cell transcriptome landscape for six major systems in human fetus

To essentially cover the whole human body, we collected 17 representative organs (esophagus, stomach, small intestine, large intestine, liver, pancreas, kidney, bladder, bronchus, lung, bone marrow, spleen, thymus, heart with artery, diaphragm, ovary and testis) in 31 different sampling positions (e.g. fundus, body and antrum of the stomach) from human fetuses at 19-21 weeks post-gestation (Figure 1A). After dissociation and non-marker-based FACS sorting, we adopted the high-precision single-cell RNA-seq method (MALBAC-DT) (Chapman et al., 2020) for library preparation and cDNA sequencing, which produced the transcriptome profile in 42,912 cells (Figure 1B). After rigorous quality control, we retained 31,208 high-quality cells, and on average each cell contained 1.9 million clean reads, 4,610 detected genes and 25,630 UMIs (Figure S1). At the same time, we also created the open chromatin landscape (Figure 1B, see below for more details). These two landscapes laid a foundation for further investigation of CGMs at both genetic and epigenetic levels (Figures 1C and 1D, see below for more details).

**Figure 1.**
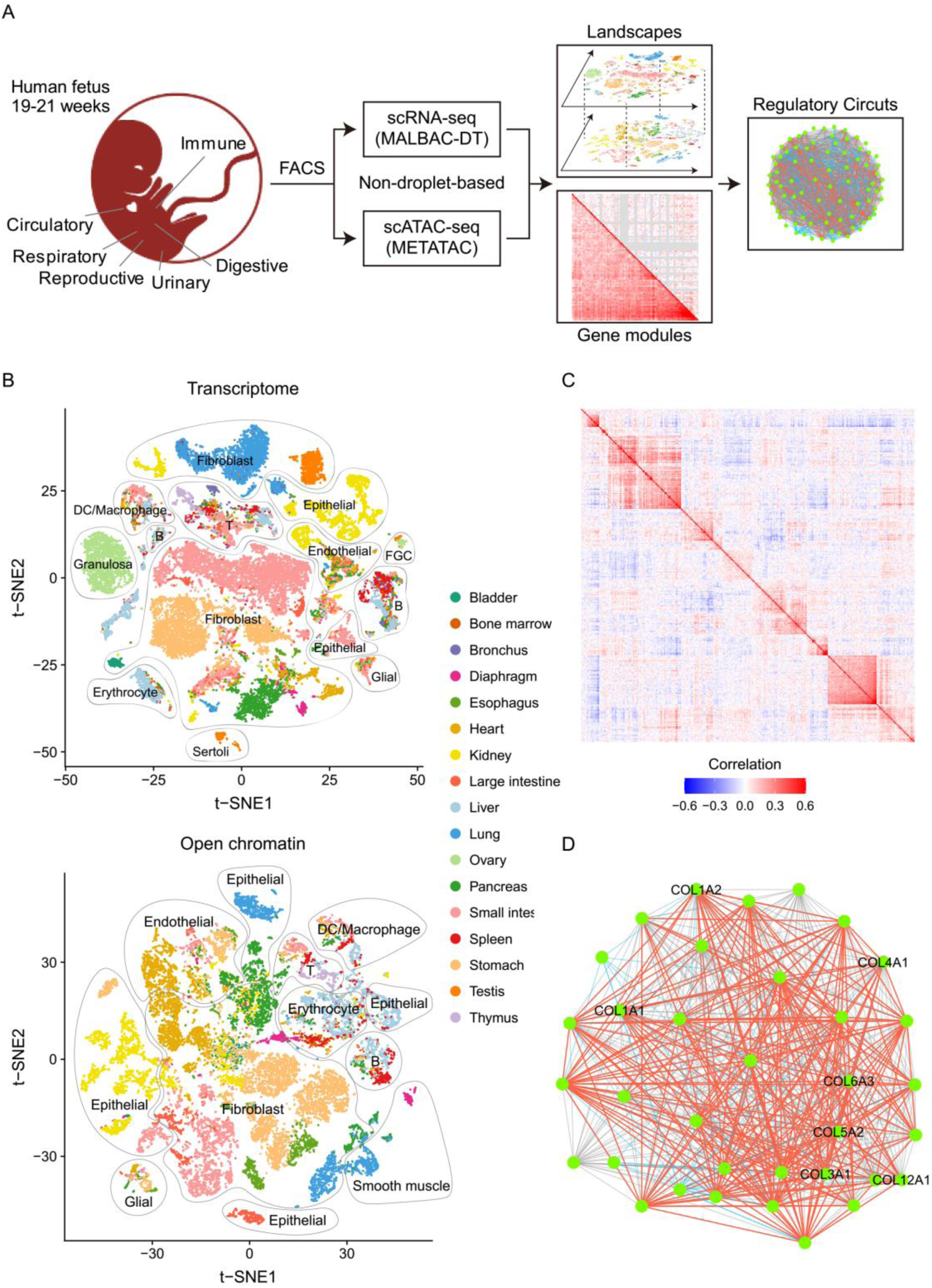
The overview of GeACT project. (A) The workflow of single-cell data production and analysis. (B) Upper panel: The *t*-SNE plot of all 31,208 cells from single-cell RNA-seq., Lower panel: The *t*-SNE plot of all 21,381 cells from single-cell ATAC-seq. Each point represents a cell, colored by organ. The clusters are annotated with the primary cell type. (C) The heatmap shows the Spearman correlation of gene expression for the CGMs genes in the stomach Fibro-ADAM28 cells. (D) Co-expression and co-accessibility network for the MD173 genes of the stomach Fibro-ADAM28 cells. Each node represents a gene. Red lines: co-expression index > 0.1 and co-accessibility index > 0.5. Blue lines: co-expression index > 0.1 only. Grey lines: co-accessibility index > 0.5 only.

To explore the cell composition of each organ, we processed the single-cell data and obtained 228 cell clusters (Table S2), each of which was annotated according to well-known marker genes from the literature (Gao et al., 2018; Li et al., 2017; MacParland et al., 2018; Pellin et al., 2019; Young et al., 2018). Then all the cells were clustered to make up the global transcriptome landscape (Figures 1B and 2A), which consisted of 6 primary cell groups common in most organs (epithelial cells, endothelial cells, fibroblasts, glial cells, immune cells and erythrocytes) (Figure 2B) as well as several cell clusters specific to sexual organs such as Granulosa cells in the ovary and Sertoli cells in the testis.

**Figure 2.**
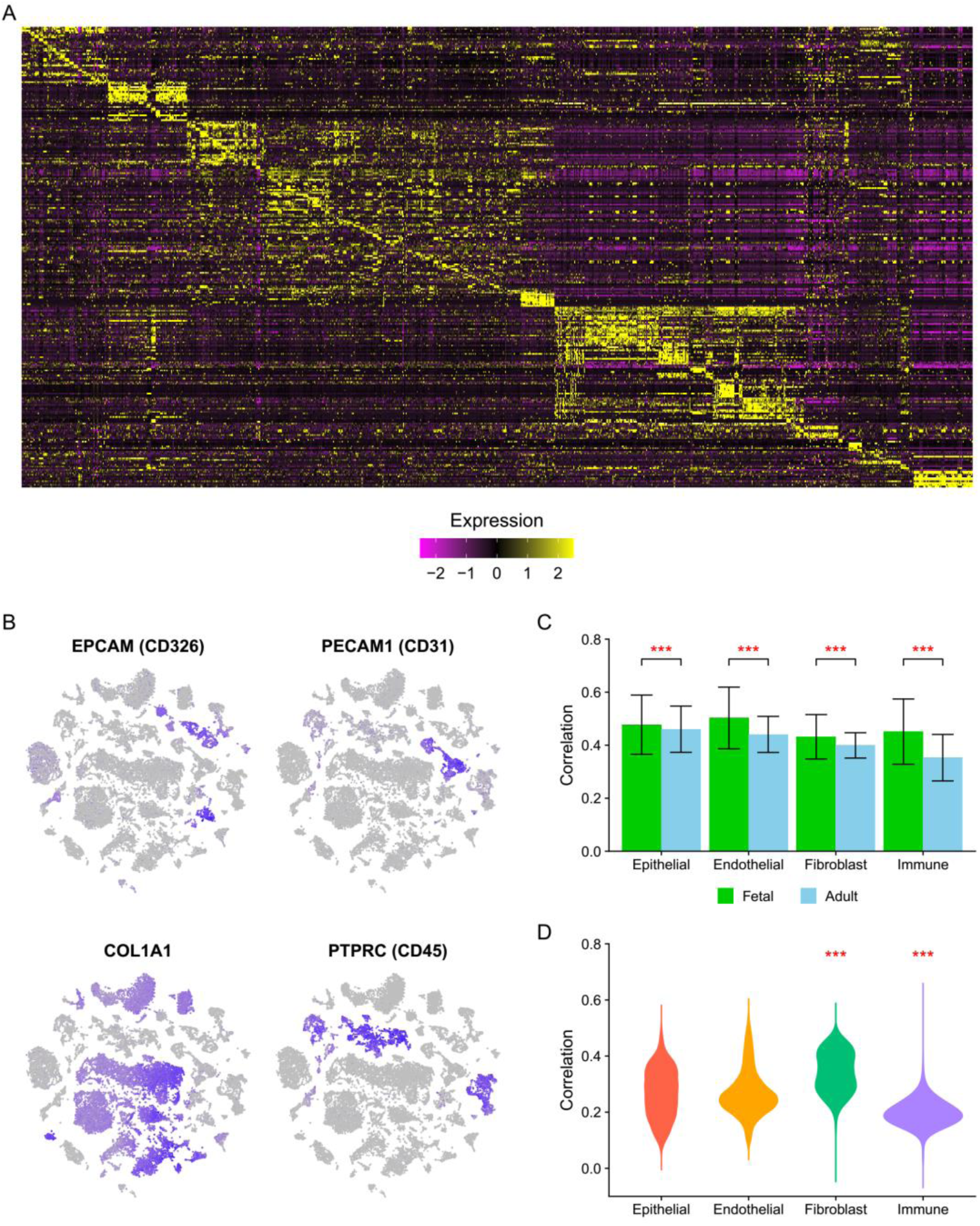
The single-cell transcriptome landscape in 17 organs. (A) The heatmap shows the signature genes in each cell type. Each row represents a signature gene and each column represents a cell. For each cell type, five cells were randomly selected for show. (B) The *t*-SNE plot of the marker genes of epithelial cells (*EPCAM*), endothelial cells (*PECAM1*), fibroblasts (*COL1A1*) and immune cells (*PTPRC*). The color changed from grey to blue as the gene expression levels increase. (C) The bar plot means the average of pairwise Spearman correlation between the cells within each organ and cell group for fetal and adult cells, respectively. The error bar means the standard derivation (Wilcoxon rank-sum tests, *** means p-value < 0.001). (D) The violin plot means the pairwise Spearman correlation between the fetal and adult cells within each organ and cell group (Wilcoxon rank-sum tests, *** means p-value < 0.001 between the corresponding cell group and any other groups).

Interestingly, compared with those in the human adult (Cao et al., 2019b; Stuart et al., 2019), the cells in the human fetus showed higher similarity within cell groups, especially for immune cells (Figure 2C). The fibroblasts showed the highest similarity between the fetal and adult stage, which indicates their unsynchronized differentiation and maturation during development and suggests the earlier development and maturation of fibroblasts than other types of cells at mid-gestation stage (Figure 2D).

As one of the organs in the digestive system, the stomach is important for food digestion and absorption (Carey et al., 1983). Different from previous work (Gao et al., 2018), which focused on the epithelial cells, we leveraged the full repertoire of stomach cells to create a single-cell landscape for the whole organ. There were 20 distinct cell types revealed in the current analysis (Figure S2A). Besides the epithelial cells (C1), where *EPCAM* was highly expressed, most cells belong to mesenchyme due to the specific expression of *VIM* (Figure S2B). Among them, we found 8 types of fibroblasts (C5-C12) according to the commonly expressed *COL1A1* but different signature genes (Figure S2C). For example, C6 was the most abundant fibroblasts where *ADAM28* was highly expressed. C12 was the proliferative fibroblasts with the high expression of cell cycle-related genes such as *TYMS*. Moreover, we found 3 types of immune cells, such as B cells (C14), dendritic cells or macrophages (C15), and T cells (C16). We also identified endothelial cells (C2), smooth muscle cells (C3 and C4), glial cells (C13), and *CACNA1A*+ cells (C17 and C18) and erythrocytes (C19). Besides the signature genes used to define cell types, transcription factors (TFs) showed distinct expression patterns across cell types (Figure S2D). For example, *FOXA2* and *EGR* were specifically expressed in epithelial cells and endothelial cells, respectively. Interestingly, *ELF3*, which played an important role in epithelial cell differentiation, was specifically expressed in both *CACNA1A*+ cells and epithelial cells, indicating that *CACNA1A*+ cells may be a group of epithelial-like cells. Based on the gene ontology (GO) enrichment analysis against signature genes, we investigated the putative functions of each cell type. Unexpectedly, different fibroblast cell types showed distinct putative functions (Figure S2E). For example, the Fibro-FBLN1 and Fibro-NRK cells were related to extracellular matrix organization but the Fibro-KCNJ8 cells were related to tube morphogenesis. Benefit from the sampling from different physiological positions, we were able to explore the spatial-specific cell-type composition. Most of the cell types showed a similar composition across different positions of the stomach. However, the body of the stomach showed a higher fraction of Fibro-FBLN1 cells but a lower fraction of visceral smooth muscle cells than the fundus and antrum (Figure S2F).

As the largest solid organ in the human body, the liver carries out many biological functions such as nutrients processing (Petersen et al., 2017) and blood storage (Brauer, 1963). In our dataset, we found 19 cell types in the liver (Figure S3A). Different from the stomach, the liver contained a substantial proportion of immune cells and erythrocytes (Figure S3B). Based on the signature genes (Figure S3C), we defined different subtypes of immune cells such as B cells (C7-C9), dendritic cells or macrophages (C10), the progenitor of Mast cells (C11), NKT cells (C12-C14) and T cells (C15). Interestingly, we found two types of erythrocytes: non-proliferative (C17) and proliferative ones (C18), which may reflect the process of blood formation. Besides several common cell types such as epithelial cells (C1), endothelial cells (C2-C4) and fibroblasts (C5), we also observed multipotent progenitors (MPPs) (C6) with the high expression of *CD34* and hepatocytes (C16) with the specific expression of *CYP3A7*. We then explored the expression of TFs across these cell types (Figure S3D). Interestingly, *MYC*, a proto-oncogene, was specifically expressed in the epithelial cells. Instead, *HMGA2* and *TFDP1* were specifically expressed in MPP cells and proliferative erythrocytes, respectively. As for the putative functions of each cell type, we were surprised to find that the Endo-DNTT cell type showed immune-related functions (Figure S3E) despite little *PTPRC* (*CD45*) expression. Although most of the cell types showed similar composition in different positions, the fraction of erythrocytes declined from segment IV to around regions (segment VII/VI/II/III) (Figure S3F), highlighting the important roles of blood supply in the segment IV (Alghamdi et al., 2017).

The kidney is an important organ in the urinary system. Although much endeavor has been made for the fetal kidney (Hochane et al., 2019; Wang et al., 2018; Young et al., 2018), the 19-20 weeks post-gestation, a key period when glomerular filtration started to significantly contribute to amniotic fluid (Rosenblum et al., 2017), was rarely covered. In the analysis of the data from the high-precision library preparation method, we were able to find 27 cell types in the kidney (Figure S4A). Different from the organs mentioned above, the kidney contained substantial epithelial cells with the specific expression of *EPCAM* (Figure S4B), some of which directly reflected the physiological structures of the kidney such as proximal tubules (C1), loop of Henle (C2 and C3), distal tubules (C4), principle cells (C5 and C6), ureter epithelium cells (C7). We also identified proliferative epithelial cells (C8). Interestingly, we found a type of EPCAM-positive podocytes (C9), which was different from the traditional one (C10) (Figure S4C). Besides epithelial cells, we also observed cap mesenchyme (C11) as well as several common cell types in other organs such as endothelial cells (C12-C14), smooth muscle cells (C15), fibroblasts (C16-C22), glial cells (C23), immune cells (C24), *CACNA1A*+ cells (C25) and erythrocytes (C26). Different cell types showed distinct expression patterns of TFs (Figure S4D). For example, *HNF4G* and *SIM2* were specifically expressed in proximal tubules and loop of Henle, respectively, but IRF6 was highly expressed in ureter epithelium cells. Moreover, two TFs in SOX family, *SOX17* and *SOX1*, were specifically expressed in endothelial cells and glial cells, respectively. Despite diverse expression across epithelial cell types, the common development-related terms indicated their common developmental stage (Figure S4E). On the other hand, several cell types showed a highly spatial-specific pattern, which may indicate the different functions across kidney positions (Figure S4F). For example, Epi-Ureter, as it was named, was specially located in Pelvis. Instead, endothelial cells were enriched in Medulla as expected.

Furthermore, the single-cell gene expression for the other 14 organs was also systematically investigated (see the website for more details). These resources made up the most comprehensive high-precision single-cell transcriptome landscape in the human for the first time.

### The architecture of gene expression profiles across organs

The comprehensive transcriptome dataset paves the way to systematically exploring the similarity of expression profiles in different organs at the single-cell resolution. Based on the hierarchical clustering of all expressed genes, cell types from different organs but with similar physiological identities (e.g. epithelial cells, endothelial cells and fibroblasts) were tended to be grouped together, suggesting their similar functions and gene expression patterns across different organs (Figure 3A). A similar pattern was also found in the clustering based on only TFs, cell surface markers or lncRNAs, which indicated that they may all contribute to the specific functions of each cell identity (Figure S5).

**Figure 3.**
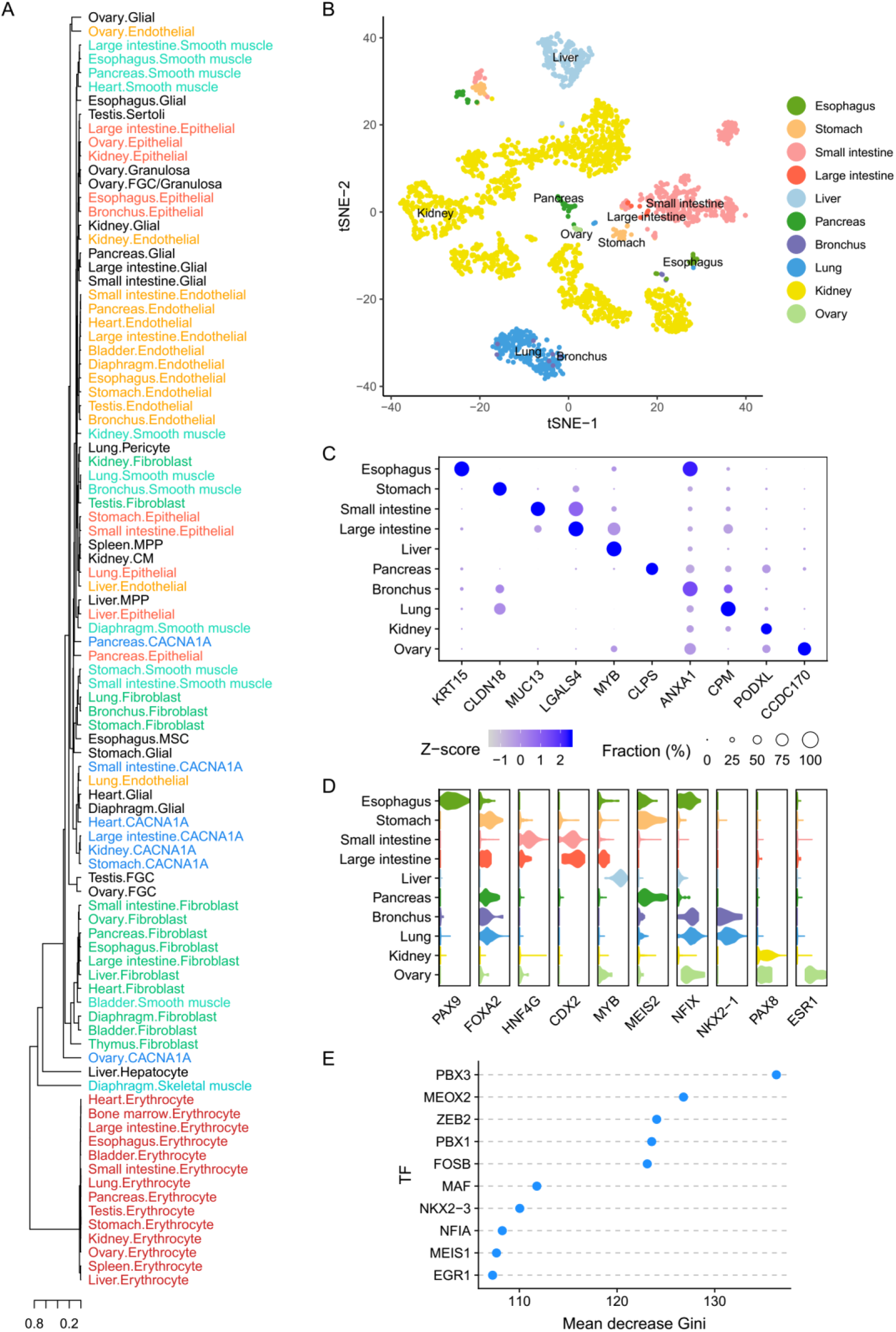
The architecture of expression profiles across cell types. (A) The hierarchy clustering plots of gene expression for non-immune cell types based on all the expressed genes, colored by cell types. (B) The t-SNE plot shows the epithelial cells in different organs. (C) The dot plot shows the signature genes of the epithelial cells in different organs. (D) The violin plot shows the signature TFs of the epithelial cells in different organs. (E) The dot plot shows the TFs classifying all cell types, ordered by mean Gini importance of the random-forest model. Only the top 10 significant TFs are shown.

On the other hand, even cells with similar identities showed distinct features across different organs. For example, the epithelial cells could be largely clustered by the corresponding organs (Figure 3B). The cells tended to be grouped with the ones from the organ in the same system, such as lung and bronchus, which indicated that this pattern was contributed by physiological differences instead of technical batch effects across organs. Several genes showed distinct expression patterns in the epithelial cells across different organs (Figure 3C). For example, the esophagus epithelial cells showed the specific expression of *KRT15*, which was reported as a signature gene of the esophagus mucosa (Mele et al., 2015). Instead, *MUC13* was specifically expressed in the epithelial cells of the small intestine and large intestine. Moreover, several TFs contributed to the differences in the epithelial cells across organs (Figure 3D). For example, *PAX9*, which played critical roles during fetal development (Mansouri et al., 1996), was specifically expressed in the epithelial cells of the esophagus. Instead, *MYB* was highly and specifically expressed in epithelial cells of the liver.

Consistent with the previous report (Tabula Muris Consortium, 2018), we found that multiple TFs significantly contribute to the variability across different cell types (Figure 3E), including *PBX3*, a key homeobox transcription factor for mesodermal commitment (Slenter et al., 2018).

### The construction of single-cell open chromatin landscape

To further explore the epigenetic mechanisms underlying the cell-type-specific gene expression profile, we isolated nuclei from 14 representative organs (except for ovary, testis, bronchus) from the corresponding fetuses. After dissociation, we used a high-precision single-cell ATAC-seq method (METATAC) for library preparation, followed by deep sequencing. In total, we captured 23,520 cells from 30 different sampling sites.

After rigorous quality control (QC) for each organ, 21,381 cells were kept for downstream analyses (Figure S1C). Averagely, each cell passed QC contained 717,814 clean reads, 79,146 unique fragments, and 38,916 detected peaks (Figure 1D and Table S3), which was much higher than previously reported mammalian tissue data (Cusanovich et al., 2018).

In order to identify cell types, we first generated 333,614 accessible chromatin regions. Then for each organ, cell types were annotated based on the cell co-embedding of the transcriptome landscape and Cicero gene activity scores (Pliner et al., 2018) using Seurat (Stuart et al., 2019), and 177 cell clusters were obtained (see Methods). The global open chromatin landscape of all cells was in accordance with the transcriptome landscape, further confirming the reliable cell typing for METATAC data (Figures 1A, 4A as well as Figures S6A and S6B). Interestingly, the genomic accessibility in TF motifs was strongly correlated with TF RNA expression levels (Figure 4B), which suggested that the chromatin accessibility could reflect TF activity veritably (Granja et al., 2019).

**Figure 4.**
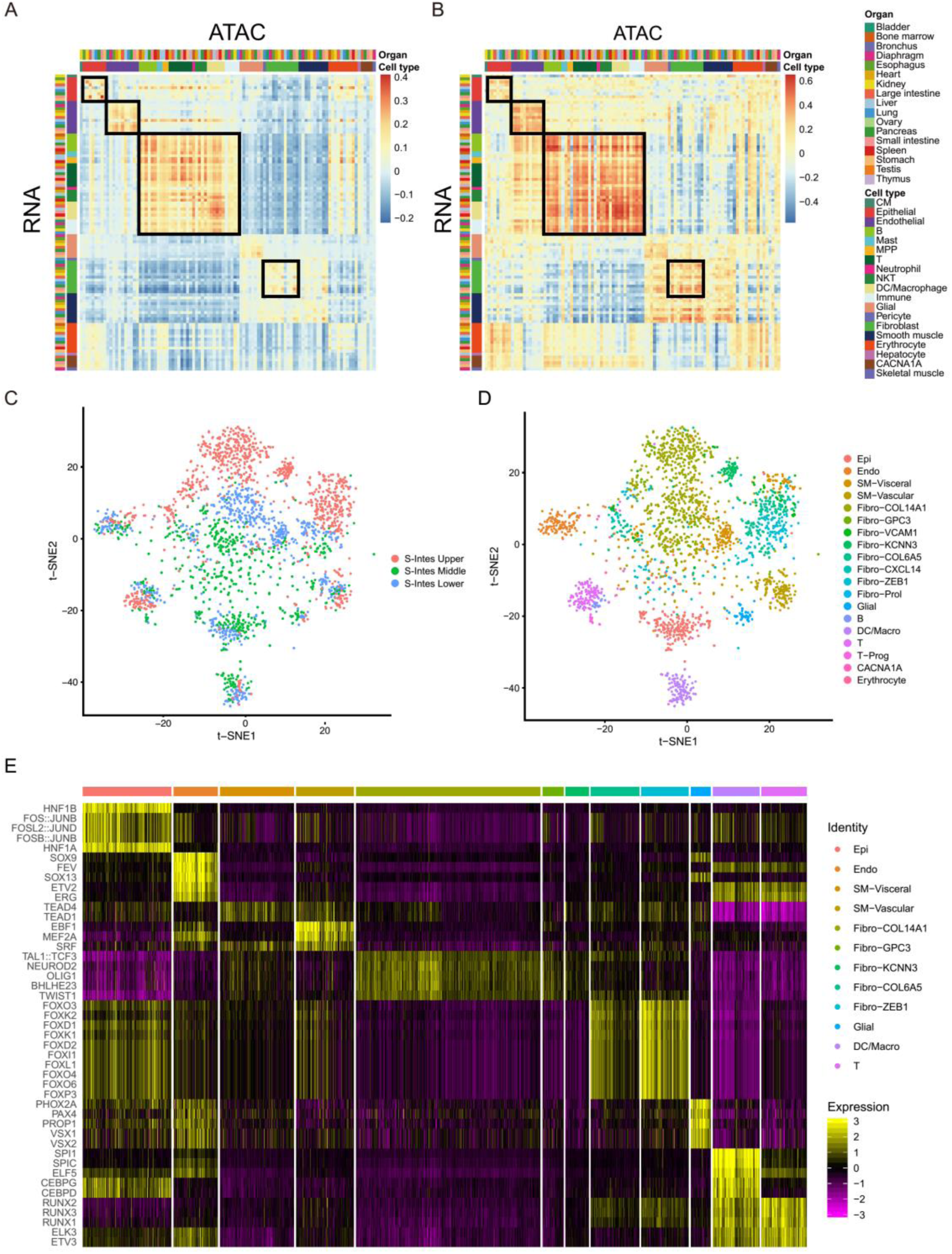
The single-cell open chromatin landscape in 14 organs. (A) Heatmap of Spearman correlations between average gene expressions and gene activity scores for common cell groups of scRNA-seq and scATAC-seq profiles. Clustered by major cell types. (B) Heatmap of Spearman correlations between average TF expressions and TF activities defined by chromVAR deviations. Clustered by major cell types. Epithelial cells, endothelial cells, immune cells and fibroblasts are highlighted. (C-E) The single-cell chromatin accessibility landscape in the small intestine. (C) The t-SNE plot shows the cells colored by sampling positions. (D) The same as (C), colored by primary cell type annotations. (E) Heatmap of marker TF motif scaled accessibilities for each cell type in the small intestine.

With the advantage of multiple sampling sites for each organ, we could compare cell type composition of different sampling sites. As an example, we showed small intestine, which was divided into upper, middle, and lower segments. We detected 19 cell types in the small intestine, including epithelial cells, endothelial cells, 4 types of immune cells, erythrocyte, 8 types of fibroblasts, glial cells, *CACNA1A*+ cells and two types of smooth muscle cells. Most cell types consisted of cells from all three parts, except for Fibro-KCNN3 and *CACNA1A*+ cells, almost all cells of which belong to the upper part. We noticed some types of fibroblasts tended to cluster according to sampling sites, such as Fibro-COL14A1 (Figures 4C and 4D).

To unravel the regulatory program underlying cell-type-specific transcriptional programs, we inferred activated TFs for each cell type based on TF motif accessibility *Z* scores (Figure 4E). Interestingly, we found that TF motifs significantly more accessible in the epithelial cells were all involved in epithelial-mesenchymal transition (EMT), like HNF1A, HNF1B, FOS and JUN family proteins. Previous research in mice showed the prevalence of epithelial cells with mesenchymal features during organogenesis (Dong et al., 2018), which revealed the mesenchymal features of epithelial cells are important for the establishment of proper organ morphology during organogenesis. For the endothelial cells, we identified SOX9, SOX13, ETV2, FEV, ERG, some of which were known essential for endothelial cell development, like SOX9 (Akiyama et al., 2004), ETV2 (Oh et al., 2015) and ERG (Birdsey et al., 2008). Two smooth muscle cell types have different marker TFs. EBF1 (Jin et al., 2014) and MEF2A (Black and Olson, 1998) binding peaks were only accessible in SM-Vascular cells but not in SM-Visceral cells, while TEAD (Liu et al., 2014) family binding peaks were more accessible in SM-Visceral cells, which may contribute to their different functions. Forkhead family motifs showed high TF *Z* scores in Fibro-COL6A5 and Fibro-ZEB1 but not in other fibroblasts, while marker TFs of Fibro-COL14A1 included neuron related TFs, such as NEUROD2 and OLIG1. In T cells, RUNX family TFs were significantly enriched, which was known to regulate T cell maturation and lineage choice (Collins et al., 2009; Egawa et al., 2007).

### The architecture of open chromatin profiles across organs

Based on the comprehensive chromatin accessibility information, we sought to explore the similarities and differences of epigenetic state across different organs with single-cell resolution. We clustered all non-immune cells based on all accessible peaks, results were highly consistent with transcriptome, which showed cells of similar epigenome but from different organs tended to cluster together (Figure 5A). Interestingly, erythrocytes from the kidney, large intestine, lung, and stomach were clustered to other cell types from the same organ instead of erythrocytes of other organs, which was different from RNA expression profiles, indicating some potential interactions with surrounding cells.

**Figure 5.**
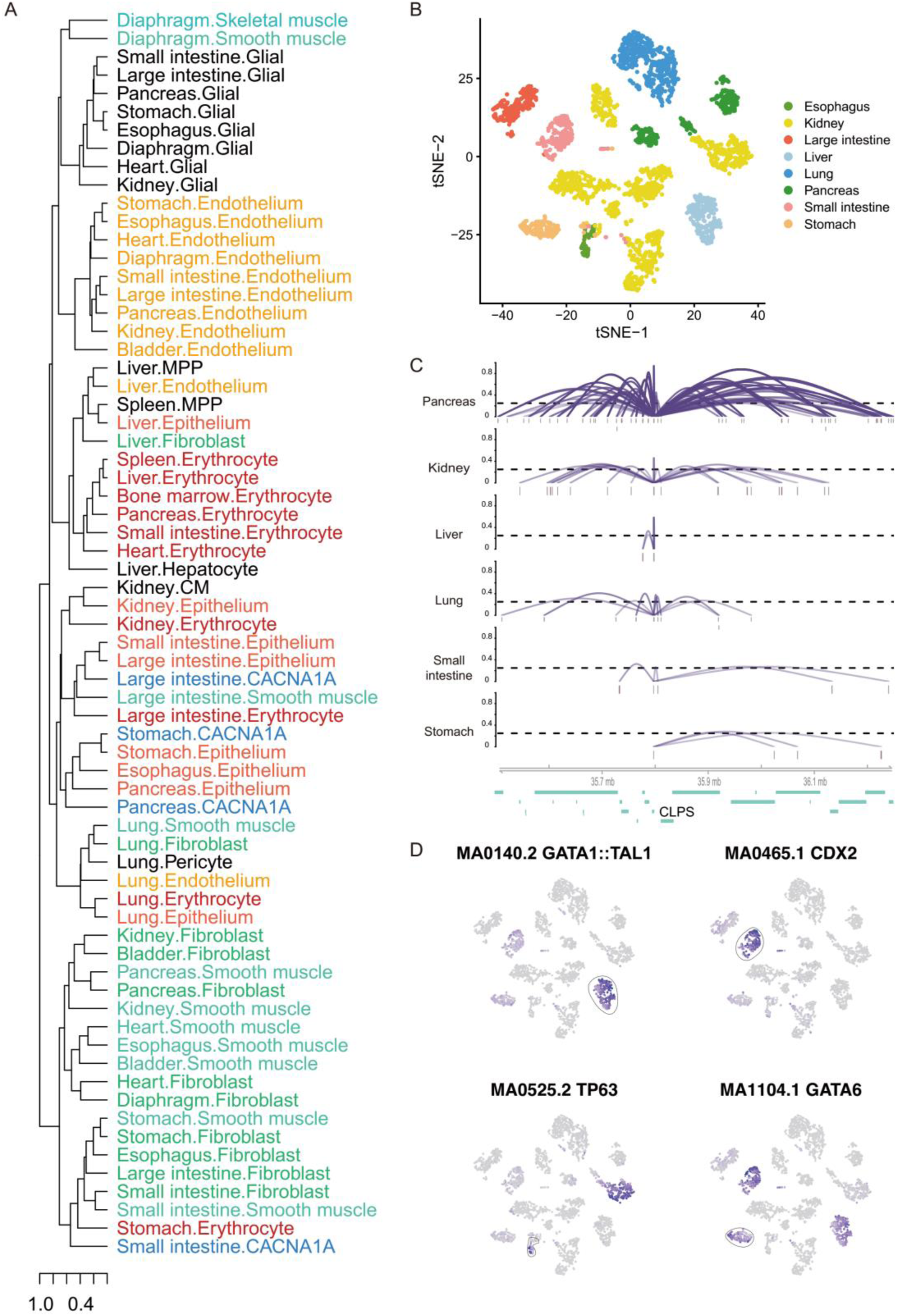
The architecture of the open chromatin profiles across cell types. (A) The hierarchy plots of open chromatin for non-immune cell types based on all accessible peaks, colored by cell types. (B) The t-SNE plot of the epithelial cells from different organs. (C) Cicero peak-to-gene connections for the CLPS locus in different organs. *CLPS* is a signature gene of epithelial cells in the pancreas. Only connections with co-accessibility score >= 0.25 are shown. (D) TF activity (defined by chromVAR deviations) overlay on single-cell ATAC t-SNE plot (as in B), showed signature TFs of the liver (GATA1-TAL1 complex), small intestine (CDX2), esophagus (TP63) and stomach (GATA6).

To characterize the overall cellular heterogeneity for epithelial, we clustered epithelial cells across diverse tissues. In accordant with RNA expression profiles, epithelial cells from the same organ largely clustered together (Figure 5B). For signature genes of epithelial cells from different organs, we associated their gene-body and promoter peaks with distal regulatory elements based on Cicero co-accessibility scores (Pliner et al., 2018), to compare the regulatory relationship across different organs. For instance, *CLPS* is specifically expressed in pancreas epithelial cells, which is a cofactor of pancreatic lipase for efficient dietary lipid hydrolysis (Borgstrom and Erlanson, 1973). The peak-to-gene connections of *CLPS* in the pancreas are much more abundant and stronger than in other organs (Figure 5C). Similar results were observed for other signature genes, such as *MUC13*, a marker gene of epithelial cells in the small intestine (Figure S6C), and *KRT15*, a marker gene of epithelial cells in the esophagus (Figure S6D). Interestingly, although *MUC13* was only expressed in epithelial cells of the small intestine at this embryonic stage (Figure 3C), the gene locus also showed strong and abundant connections in epithelial cells of the pancreas, esophagus, stomach and large intestine. Previous research revealed that *MUC13* is a cell surface glycoprotein highly expressed in epithelial tissues of gastrointestinal and respiratory tracts (Williams et al., 2001), and is a potential pancreatic cancer diagnostic marker (Khan et al., 2018). The open chromatin profiles indicate the regulatory potential for future expression in these organs.

Based on TF motif accessibility *Z* scores (Schep et al., 2017), we inferred TFs that regulate the distinguishable expression profiles (Figure 5D). GATA1-TAL1 complex showed specific activity in liver epithelial cells. CDX2 exhibited high activity in small intestine epithelial cells, but not in large intestine epithelial cells, though it is highly expressed in both cell groups (Figure 3D). TP63 was active in both esophagus epithelial cells and renal pelvis Epi-Ureter cells. GATA6 showed high activity in epithelial cells of the liver, small intestine and stomach.

### The correlated gene module and the integrative regulatory circuit

The high-precision data offered a great chance to delineate correlated gene modules (CGMs) across cell types (Chapman et al., 2020; Chihara et al., 2018). For better robustness, we selected 10 cell types with the highest numbers of cells analyzed for CGM detection (see Figures S7A-D for more details) and obtained 227 non-redundant CGMs with the gene number in each CGM from 10 to 240 (Table S4). Each CGM showed distinct correlation profiles across cell types (Figure 6A). Interestingly, more than 60% of CGMs showed enriched TFs, which reflected on the contribution of TFs on the regulation of co-expressed genes. The enriched protein-protein interactions (PPIs) were observed in half of the CGMs, which indicated that the correlated transcription was a key process for the synchronization of protein activities. On the other hand, 69.2% and 47.6% of CGMs contained enriched GO terms and KEGG pathways, which indicated the similar biological functions of correlated genes. Although protein-coding genes constitute the majority (more than 90%) of CGMs, there were substantial non-coding genes in each CGM (Figure 6B), which indicated the similar functions of correlated genes with different gene types. Unexpectedly, for most of the CGMs, genes were scattered in different chromosomes, expect two CGMs made up of mitochondrial genes (Figure 6C), which indicated that correlated genes are merely connected by genomic proximity (i.e. *cis*-effect). Instead, the CGMs with higher correlation were more likely to contain common upstream TF regulators, which indicated *trans*-effect was the primary driving force for correlated genes. In addition, high-correlation CGMs tended to contain enriched PPIs, GO biological process and KEGG pathways, which further highlighted the collaborative mode in the functioning of genes (Figure 6D).

**Figure 6.**
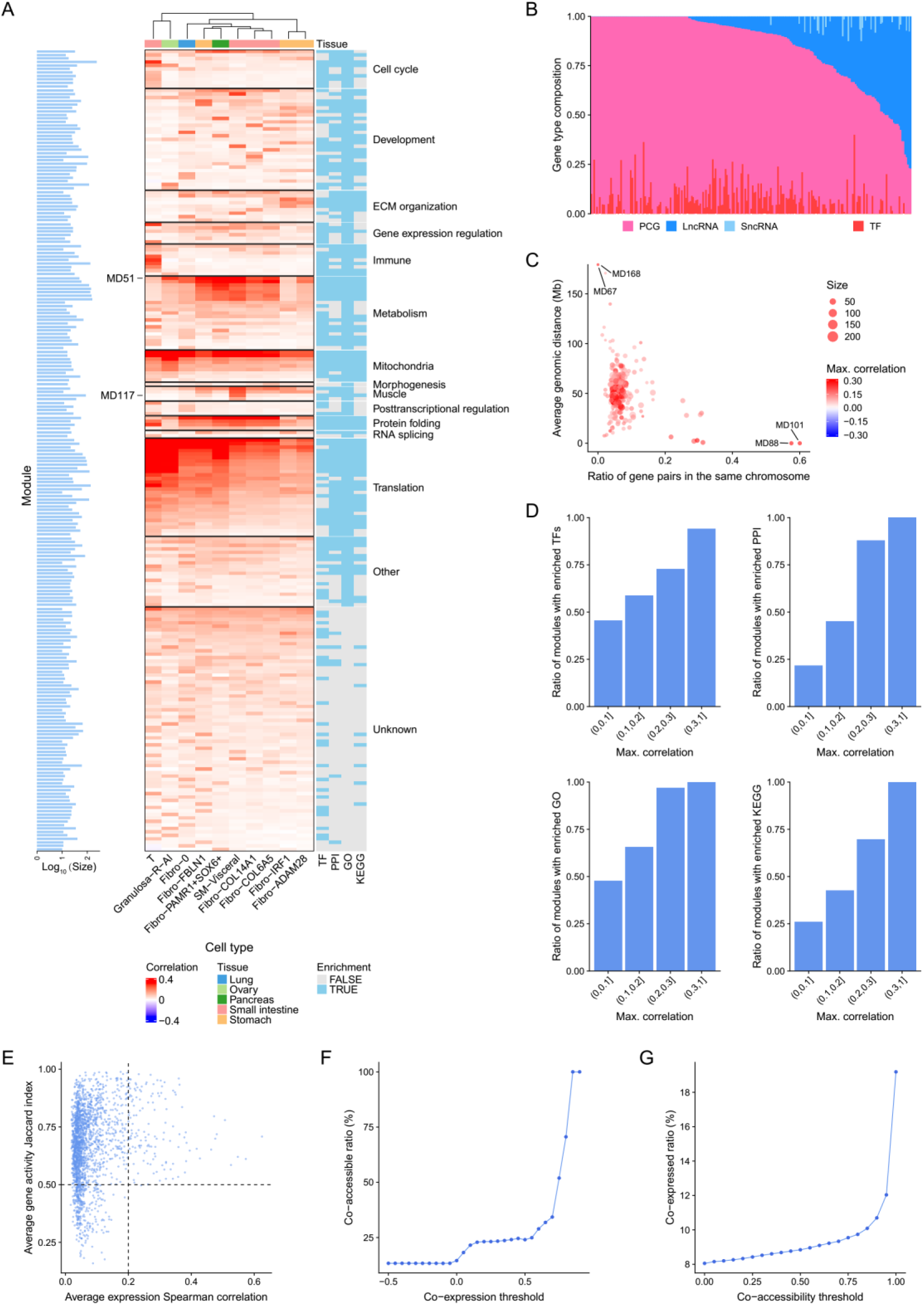
The co-expression gene module map across cell types. (A) Left: the bar plot shows the number of genes in each gene module. Middle: the heatmap shows the average Spearman correlation of gene expression levels in each gene module (row, n=227) and each cell type (column, n=10). Right: the heatmap shows whether the genes in a gene module have enriched TF or miRNA regulators, gene ontology (GO) terms, KEGG pathways and protein-protein interactions (Hypergeometric test, FDR < 0.05). Gene modules were grouped by similar biological functions and then sorted by average correlation. (B) The bar plot shows the gene type composition in each gene module, ordered by the fraction of protein-coding genes. PCG: protein-coding gene. (C) The scatter plot shows the genomic distribution pattern of genes in each gene module. The x-axis means the ratio of gene pairs in the same chromosome and the y-axis means the average genomic distances of genes. The maximum correlation means the maximum value of the average of gene-gene correlation for each cell type. (D) The ratio of gene modules with enriched TF, protein-protein interaction (PPI), GO biological process and KEGG pathway for gene modules with different maximum correlation scores. (E) The scatterplot comparing the co-expression index and the co-accessibility index of each CGM (n=225, two mitochondria-related CGMs: MD88 and MD101, were excluded because these genes aren’t located in the nuclear genome) and each cell type (n=9). 225*9 = 2025 points in total. The co-expression index was defined by the average Spearman correlation coefficients of the expression level each pair of genes within CGM, and the co-accessibility index was defined by the average Jaccard indexes of binary gene activity scores of each pair of genes (F) The line chart shows the ratio of highly co-accessible (co-accessible index >= 0.98) gene pairs changes over the co-expression threshold. (G) The line chart shows the ratio of highly co-expressed (co-expression index >= 0.2) gene pairs changes over the co-accessibility threshold.

A CGM may show different co-expression levels in different cell types (Figure 6A). We assumed that if genes are highly co-expressed in a cell type, the epigenetic state of the regulatory genomic elements of these genes should change synchronously in this cell type. To verify this hypothesis, we quantified the co-accessibility of two genes using the Jaccard index of binary gene activity scores of METATAC data calculated by Cicero (Pliner et al., 2018). For 9 of the 10 cell types with more than 500 cells in RNA expression profiles (except for ovary Granulosa-R-Al cell type due to the lack of open chromatin profile of ovary), we calculated the average RNA expression Spearman correlation coefficients and average ATAC gene activity Jaccard index of all pairs of genes within each CGM (Figure 6E), denoted as co-expression index and co-accessibility index, respectively. In many cases a CGM with high co-accessibility showed low co-expression in a cell type, however, almost all CGMs with co-expression index higher than 0.2 have co-accessibility index higher than 0.5 in the corresponding cell type. We next compared the co-expression and co-accessibility of each gene pair within the same CGM for each of the 9 cell types, by setting different co-expression threshold to analyze the ratio of highly co-accessible pairs, and vice versa. The ratio of highly co-accessible pairs increases as the co-expression threshold (Figure 6F) and reaches 100% when the co-expression threshold is larger than 0.8. The ratio of highly co-expressed gene pairs also increases as the co-accessibility threshold, however, even for totally co-accessible pairs, only less than 20% are highly co-expressed (Figure 6G). These results indicated that the co-accessibility of gene regulatory regions is a necessary but insufficient condition for co-expression of a pair of genes.

The CGMs ubiquitous in multiple cell types were usually involved in the basic biological processes such as metabolism, protein folding and translation. For example, MD51, a CGM with 48 genes, was highly correlated in all the 10 cell types such as Fibro-FBLN1 in the stomach and Fibro-PAMR1+SOX6+ in the pancreas (Figures 7A and 7B). There are many known functionally similar proteins included in this module, such as 4 heat shock protein chaperons, 4 phosphatases, 4 Activator Protein-1 (AP-1) TFs, 3 transcription initiation factors, 2 GTPase, 2 NF-kB inhibitors and so on. The enrichment analysis showed that MD51 was related to stress response and enriched in the MAPK signaling pathway (Figure 7C). Moreover, the protein products of the genes in MD51 such as FOS and JUN formed a complex (Figure 7D) related to stimulation response, which was important for proliferation and differentiation (Angel and Karin, 1991; Cook et al., 1999). Similar to expression correlation, the genes in MD51 showed highly accessibility correlation in both cell types (Figures 7F) and the similar difference between the two cell types (Figure S7E). To unravel the TF regulators, we performed motif enrichment analysis in the regulatory regions of these genes, defined by Cicero co-accessibility (see Methods), and then calculated the average co-expression coefficient with these genes for each TF. Six TFs showed both significant motif enrichment within the regulatory regions and high co-expression with these genes (Figure 7E). EGR family, C2H2-type zinc-finger TFs, such as EGR1, EGR2 and EGR3 were identified. EGR1 was involved in stress response under disease condition (Ponti et al., 2015; Stuart et al., 2005), EGR2 was reported to suppress the c-Jun NH2-terminal protein kinase (JNK)-c-Jun pathway, and EGR3 was an immediate-early growth response gene which is induced by mitogenic stimulation (Patwardhan et al., 1991). Both EGR2 and EGR3 played vital roles in the immune system (Taefehshokr et al., 2017). IRF1 displayed a remarkable function in the regulation of cellular responses (Kroger et al., 2002).

**Figure 7.**
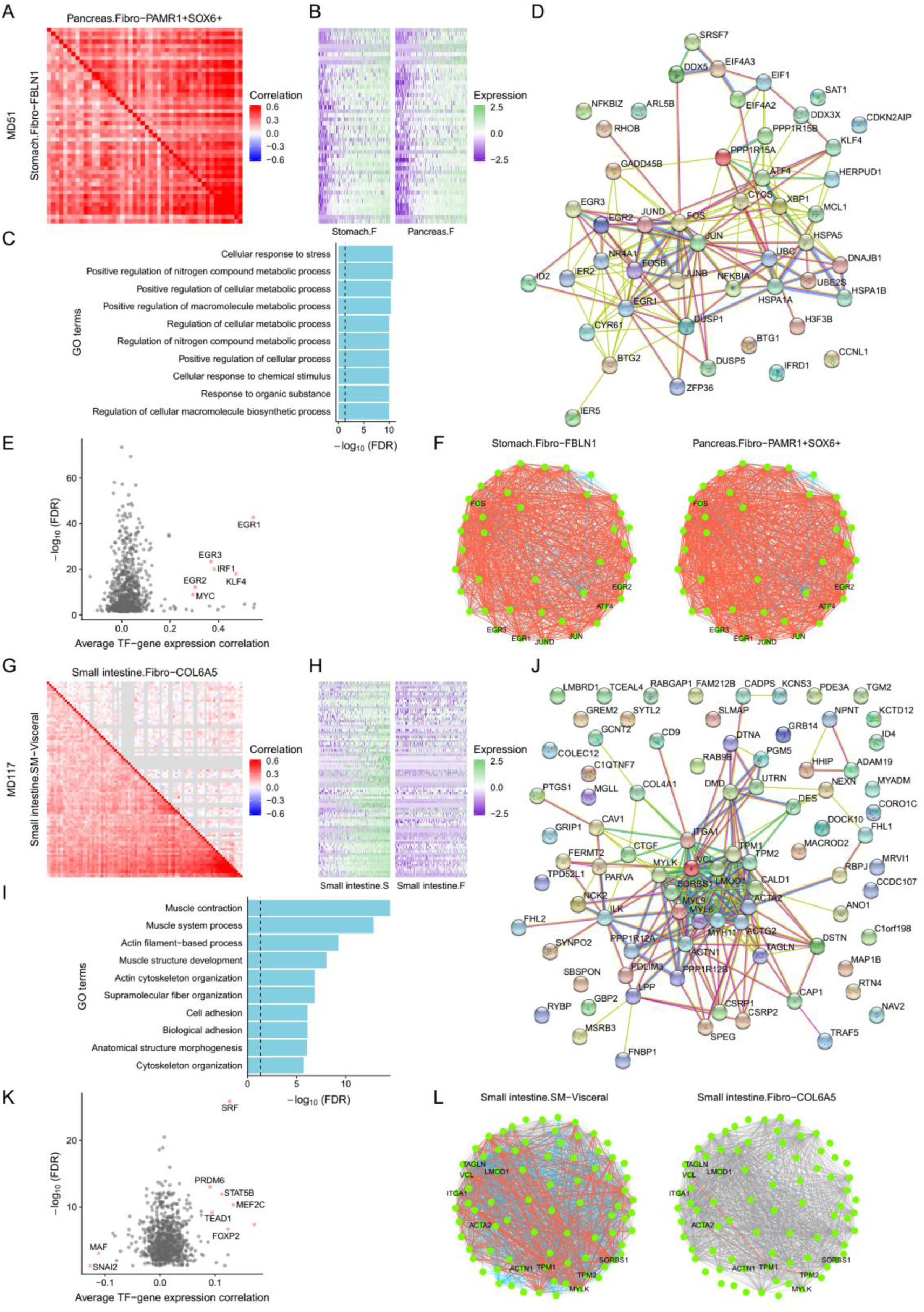
Representative gene modules. Two gene modules are shown: MD51 (A-F) and MD117 (G-L). (A and G) The heatmaps show the Spearman correlation of gene pairs. The grey blocks mean the correlation of genes expressed in less than 10% cells of the corresponding cell type. (B and H) The heatmaps show the relative gene expression levels for each gene (row, ordered as A and G, respectively) in each cell (column, only 500 randomly sampled cells are shown, ordered by expression level). (C and I) The bar plots show the enriched terms for enriched GO terms. Only the top 10 significant terms are shown. (D and J) The protein-protein interaction for the genes in the specific gene module. The blue, purple and yellow lines mean the PPIs are supported by curated databases, experimental determination and text mining, respectively. (E and K) The scatter plots show the TF motif enrichment in regulatory regions (see Methods for more details) of CGM genes (x-axis) and the average Spearman correlation coefficients of TF with CGM genes in the pancreas Fibro-PAMR1+SOX6+ cells (E) and small intestine SM-Visceral cells (K). The TFs with the FDR < 1e^-5^ and the average co-expression > 0.2 (E) < −0.1 or > 0.09 (K) are highlighted. (F and L) Co-expression and co-accessibility network of MD51 genes and MD117 genes, respectively. Each node represents a gene. Red lines: co-expression index > 0.25 and co-accessibility index > 0.9. Blue lines: co-expression index > 0.25 only. Grey lines: co-accessibility index > 0.9 only.

Meanwhile, the cell-type-specific CGMs usually reflected the function of the corresponding cell type. For example, MD117, a CGM contained 87 genes, was shown to be correlated only in SM-Visceral cells in the small intestine (Figures 7G and 7H). The enrichment analysis showed many smooth muscle-related functions (Figure 7I). In the protein level, the genes in MD117 formed the actin and myosin (Figure 7J), which was important for muscle contraction (Sweeney and Hammers, 2018). Interestingly, although expression correlation showed a highly cell-type-specific pattern, the co-accessibility patterns of MD117 genes were similar between SM-Visceral cells and Fibro-COL6A5 cells in the small intestine (Figure 7L and S7F). Based on the open chromatin data, we identified 5 regulatory TFs that were positively correlated with these genes and significantly enriched in their regulatory regions in SM-Visceral cells (Figure 7K). SRF and MEF2C were known essential TFs for myogenesis, and important in maintaining the differentiated state of muscle cells (Black and Olson, 1998; Miano, 2003). PRDM6 was involved in the regulation of vascular smooth muscle cell (VSMC) contractile proteins, suppression of differentiation and maintenance of the proliferative potential of VSMC (Davis et al., 2006). Inhibition of STAT-5B suppressed thrombin-induced VSMC growth and motility (Cao et al., 2006). RPBJ, the major mediator of Notch signaling, was important for maintaining muscle progenitor cells and generating satellite cells (Vasyutina et al., 2007). TEAD1 played an important role in inhibiting smooth-muscle specific gene expression by competing with myocardin binding to *SRF* (Liu et al., 2014). Besides, TFs anti-correlated with module genes, such as SNAI2 and MAF, although without significant motif enrichment, were also functionally related to this CGM. SNAI2 acts as a transcriptional repressor to prevent the occupancy of MYOD on myogenic differentiation-specific regulatory elements (Soleimani et al., 2012). MAF was a leucine zipper-containing TF acting as a transcriptional activator or repressor, and was up-regulated during myogenesis through MYOD (Serria et al., 2003) (Figure 7L). Different from the pattern in MD117, the co-expression and co-accessibility patterns in MD34 were both quite different between different cell types (Figures S7G-I), further highlighting the effect of open chromatin stages on the correlated gene expression levels within CGMs.

## Discussion

In the mid-gestation, the human fetus undergoes massive organ development and maturation. Our high-precision single-cell omics data identified over 200 distinct types of cells in all six major systems. Each cell type presents unique gene expression patterns, chromatin states as well as biological functions. Comparative analysis on epithelial cells among distinct organs showed that, while harboring similar marker genes, these cell types presents organ-specific gene/TF-expression patterns, implying that these critical molecules potentially regulate the organ-specific functions at their microenvironments.

Genes with high inter-tissue expression correlation usually shared similar upstream regulators or similar functions (Segal et al., 2004). However, little was known for correlated gene module (CGM) profiles within cell types during fetal development (Chapman et al., 2020; Chihara et al., 2018). We, for the first time, delineate core CGMs and underlying circuits based on the unbiased, high-precision omics data across multiple fetal organs. The 227 identified CGMs from ten cell types largely enriched potential functional TFs. Of note, the tissue/cell-type-specific CGMs showed clear transcription factor-based gene regulatory networks among the known and unknown regulon genes.

With the advantage of our high-precision single-cell transcriptome and open chromatin data, we could further combine co-accessible peaks’ motif enrichment and transcription factor-gene co-expression information to reveal functional TFs regulating each CGM (Figure 7E). Meanwhile, we show that co-accessibility is a necessary but not sufficient condition for co-expression (Figures 6F and 6G). The sophisticated symphony-like coordination between the epigenetic chromatin status and gene transcription we revealed could contribute to the effective regulation of cell-type-specific functions, as well as the establishment and maintenance of cell identity during development.

It is evident that GeACT has provided much needed and high-precision dataset as well as novel insights much beyond cell typing. We anticipate that GeACT, when expanding to all human tissues, normal or diseased, will eventually provide the understanding of the human functional genome.

## Acknowledgments

This work was made possible by support from Beijing Advanced Innovation Center for Genomics (ICG) at Peking University. The sequencing experiments were performed at Peking University High-Throughput Sequencing Center, with assistance from Chenyang Geng, Yun Zhang, Jing Sun, Yang Xu and others staffs. Part of the data analysis was performed on the Computing Platform of the Center for Life Sciences of Peking University, with help from Fangjin Chen, Ting Fang and Wenzhong Zhang on server management. We thank Yan Chen for FACS technical assistance, and Zhidan Song for assistance in establishing MALBAC-DT automated workflow. Dr. Yiqin Gao and his lab members provide helpful comments during discussion.

## Author Contributions

X.S.X., G.G., F.Tang, and J.Q. conceived the project. X.S.X., W.M., A.R.C., D.F.L., and Y.H. developed the MALBAC-DT technique and established the automated workflow. X.S.X., H.W., L.T., and D.X. developed the METATAC technique. M.Y., L.Y., J.Y., Y.M., Y.G., K.C., Y.Z., and X.Liu performed sample preprocessing. F.Z., W.M., H.W., G.Y., Jing W., and R.H. performed cell sorting. W.M., G.Y., and Jing W. performed MALBAC-DT experiments. H.W. and Y.A. performed METATAC experiments. F.Tian performed MALBAC-DT data analysis with help from A.R.C., Z.C., and L.W.. X.Li performed METATAC data analysis with help from W.S.. S.L. and D.Y. developed the website with help from F.Tian. S.Z. helped the maintenance of the computer infrastructure. F.Tian, F.Z., X.Li, W.M., and H.W. wrote the original manuscript with input from all authors. X.S.X, G.G., F.Tang, A.R.C., L.T., D.X., and W.S. reviewed and edited the manuscript.

## STAR★Methods

### KEY RESOURCES TABLE

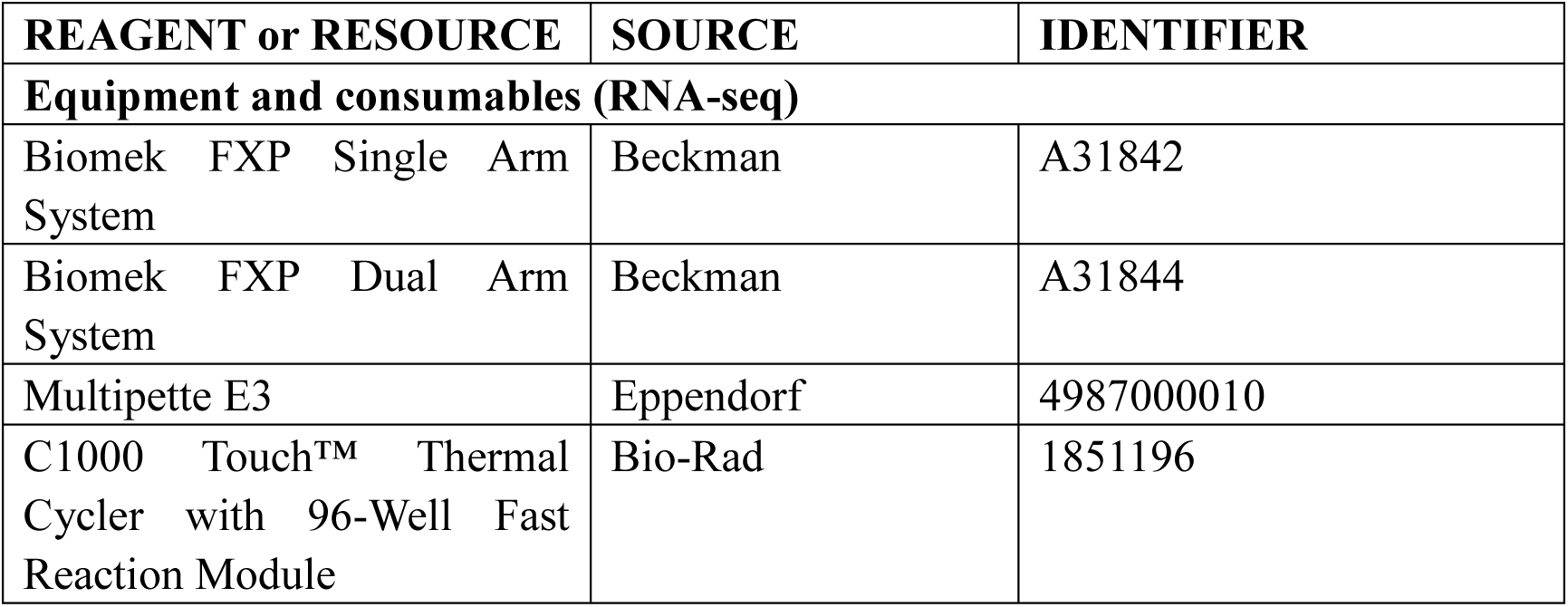
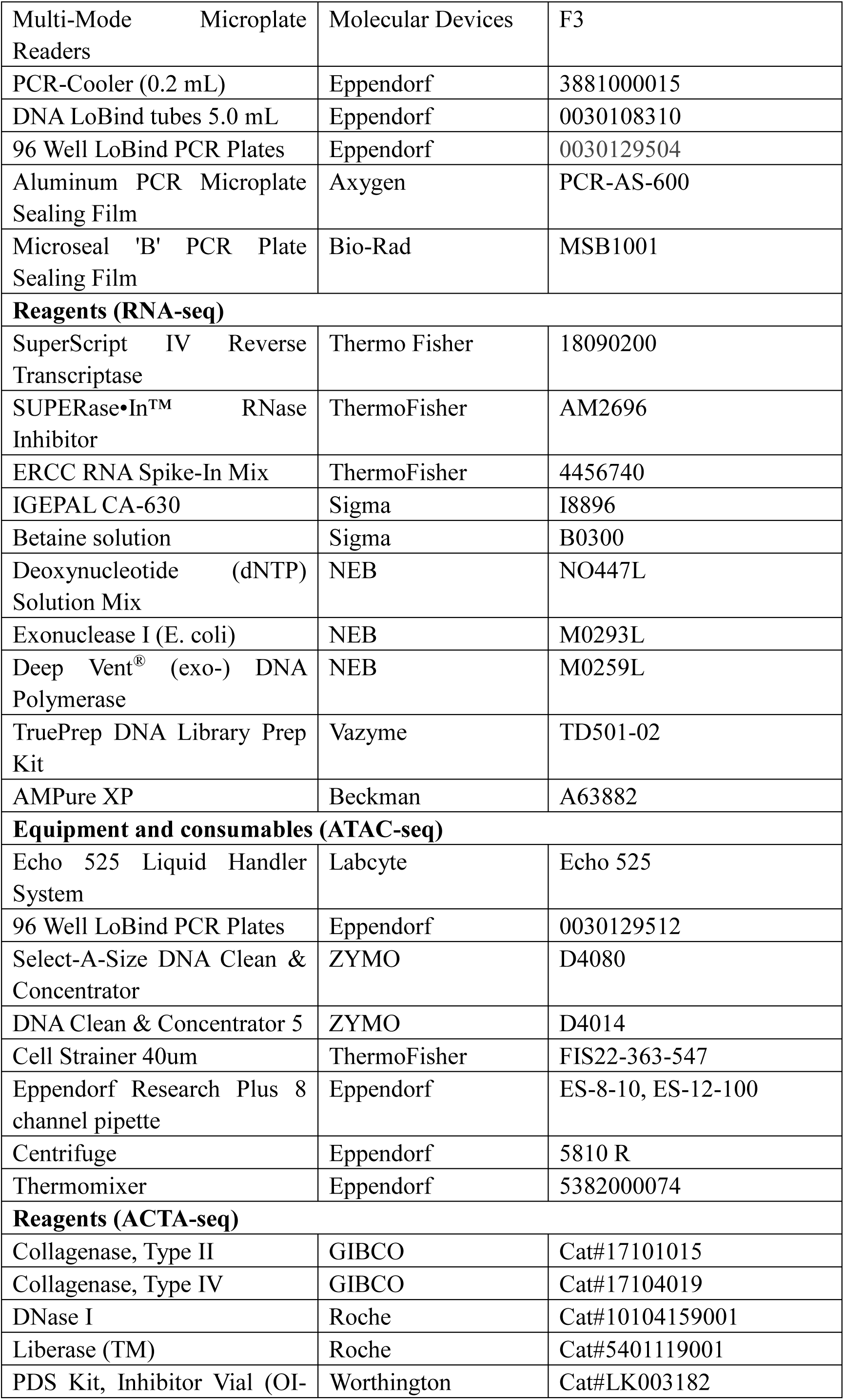
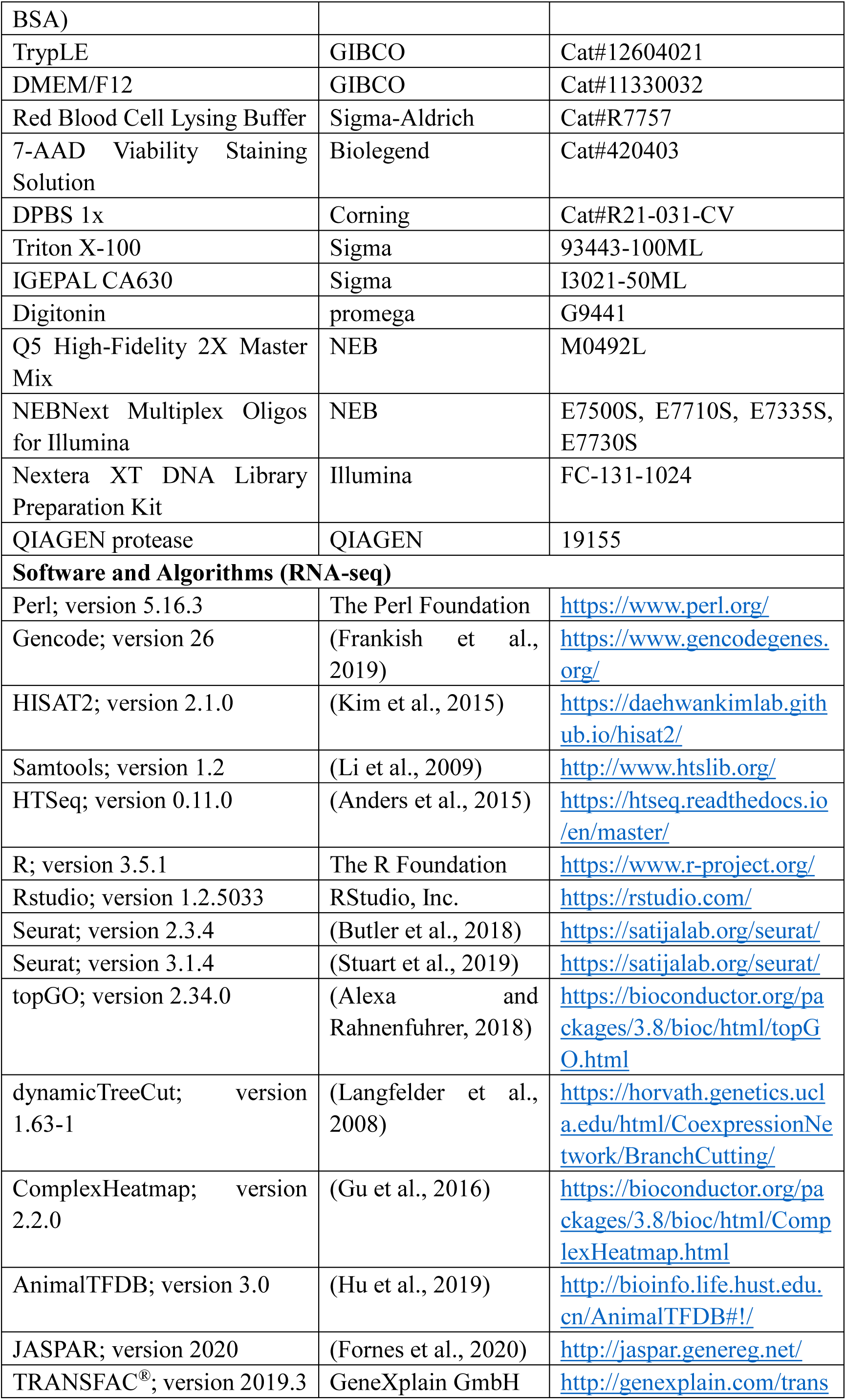
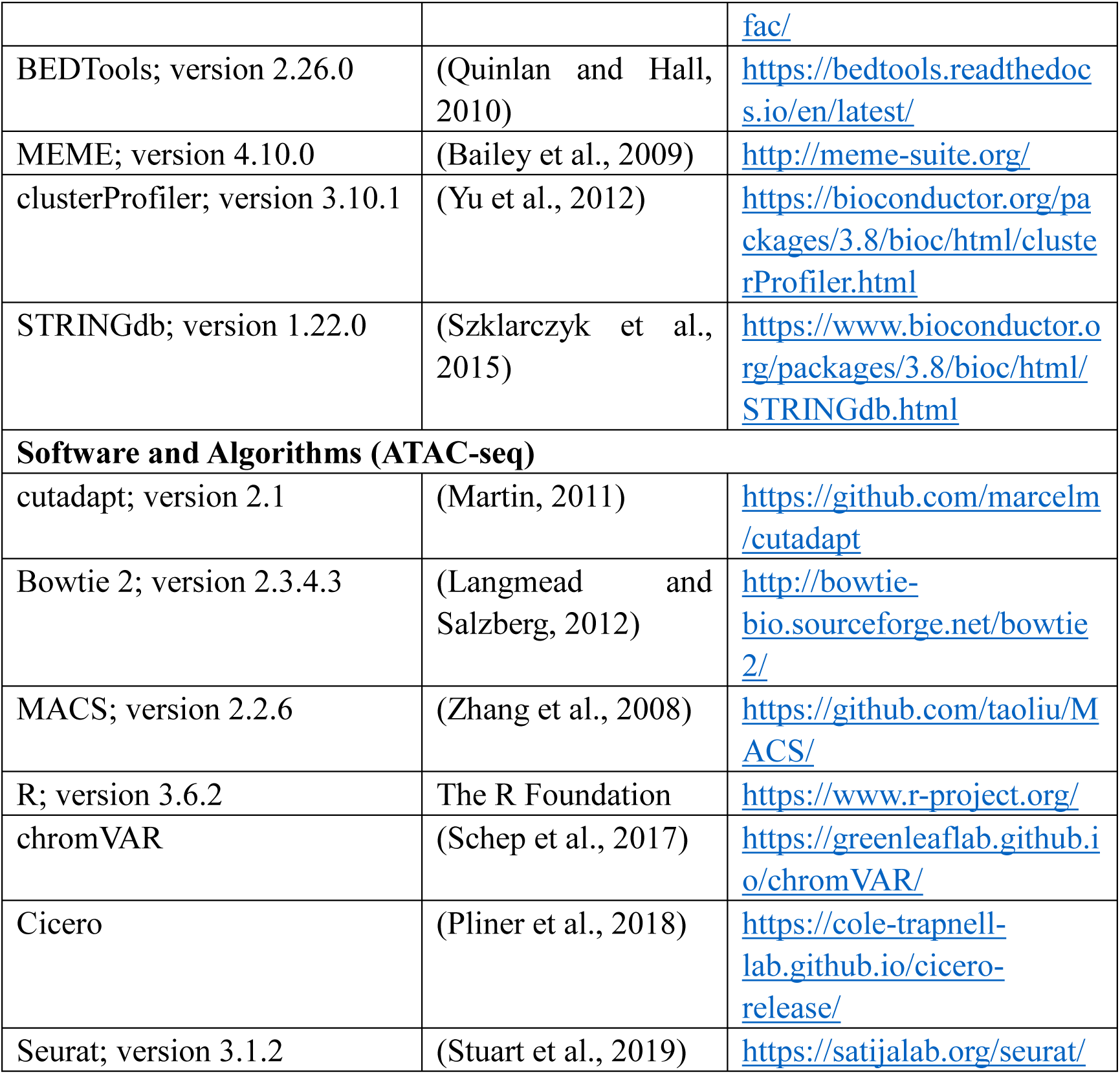

### LEAD CONTACT AND MATERIALS AVAILABILITY

Further information and requests for reagents may be directed to, and will be fulfilled by, the Lead Contact, X.S.X.

### EXPERIMENTAL MODEL AND SUBJECT DETAILS

#### Human tissues

This study was approved by the Reproductive Medicine Ethics Committee of Peking University Third Hospital (Research License 2019SZ-004). The pregnant donors underwent medical termination of pregnancy due to conditions such as cervical insufficiency, infection, eclampsia, inevitable abortion, etc. All the patients voluntarily donated the fetal tissues and signed the detailed forms of informed consent.

### METHOD DETAILS

#### Sample Dissection and Single-cell Isolation

Tissues were immediately processed to the single-cell dissociation after specimen resection. 17 organs were included in the study and the protocols of individual organs are described below.

#### Bladder

The urothelium was detached from the bladder muscle and washed twice with resuspension buffer (DMEM + 10% FBS). Then it was minced with dissecting scissors, followed by digestion at 37℃, 1000rpm sequentially in digestion buffer (2mg/ml collagenase II + collagenase IV in DMEM) for 25min and TrypLE for 5min. Cells were subsequently filtered through a 40um strainer. After wash, cells were collected by centrifugation and then stained with 1:40 7-AAD before sorting.

#### Bone marrow

Bones that excised were firstly rinsed in DMEM/F12 with 10% FBS. The bone marrow cells were flushed out by a 10ml syringe containing DMEM/F12 complemented with 10% FBS. The collagenase II/IV at 2.5mg/ml was then used to flush the bone marrow cells again. The aspirated cells were gone through a 40µm filter, centrifuged at 300g for 10 minutes. After carefully removed the supernatant, the cells were resuspended in 3ml PBS and incubated with 15ml of ACK lysis buffer for 3min at room temperature to remove the red blood cells twice. In order to excluded nonviable cells, 7-AAD was used before the FACS analysis.

#### Bronchus

Bronchus was divided into two parts, main bronchi 1-6 and main bronchi 7-12. Tissues were washed by DMEM containing 10% FBS, and then transferred into tubes containing 1 mL of papain (50 μg/ml). The tubes were incubated at 37℃ for one hour and twenty minutes with shaking at 1000 rpm. We pipetted up and down every 5 minutes to accelerate the process. After digestion, samples were filtered through a 40-μm nylon cell strainer, and then centrifuged. The cell pellets were resuspended in DMEM (contained 10% FBS), centrifuged at 300g for 5 min. The supernatant was removed, 5 μl 7-AAD and 200 μl PBS (plus 0.1% BSA) were added to the cell pellets. After incubation at RT for 10 min in a dark place, the cell suspension was mixed with a certain amount of PBS (plus 0.1% BSA), depending on the cell number and ready for FACS.

#### Diaphragm

The diaphragm was dissected, washed and minced in the digestion buffer (2mg/ml collagenase II + collagenase IV in DMEM). After that, tissue pieces were digested in a thermomixer at 37℃, 1000rpm for 25min and filtered through 40μm strainer. The collected cells were then washed twice with resuspension buffer (DMEM + 10% FBS), centrifuged and resuspended in 0.1% BSA with 1:40 7-AAD. Following filtration, single living cell was sorted into the well of 96-well plate with FACS.

#### Esophagus

Esophagus was firstly washed in DMEM which containing 10% FBS. It was then transferred to a tube and minced with the scissor. After mechanically dissociation, 1.5ml of 2.5mg/ml collagenase II/ IV mixture (GIBCO, 17101015, 17104019), 0.1 mg/ml DNase I (Roche, 10104159001) were added. The tube was incubated on a shaker at 37℃ for further dissociation. After about 45 minutes, the isolated cells were collected by certification (800g, 5min) and subsequently washed once in DMEM/F12 with 10% FBS. Cells were filtered through 40μm strainer, pelleted again, and resuspended in 200μl PBS (contained 0.1%BSA) with 5μl 7-AAD for dead cell exclusion. After 10 minutes for incubation in the dark, the cells were finally resuspended in FACS buffer waiting for cell sorting.

#### Heart

The sample covered four main zones (Left Atrium, Left Ventricle, Right Atrium, Right Ventricle) and two valves (Left and Right). Besides, we also separated interventricular and aorta under a microscope. Tissues were washed with DMEM containing 10% FBS and cut into pieces. Tissues were digested into single-cell suspension with 1 mL of collagenase Ⅱ/collagenase Ⅳ (2.5 mg/ml) and DNaseⅠ (0.1 mg/ml) at 37℃ for ten minutes with shaking at 1000 rpm. We used a 40-μm nylon cell strainer to filter the digested tissues, follow by centrifugation at 800g for 5min. DMEM containing 10% FBS was added to the cell pellets and the cell suspension was centrifuged again. After removal of the supernatant, cell pellets were mixed with 5 μl dye and 100 μl PBS (plus 0.1 %BSA) and then incubated at RT for 10 min in a dark place. According to cell number, PBS (plus 0.1 %BSA) was added to the cell suspension. All heart tissues were treated the same way, except aorta needed a longer time than others due to harder dissection and less cells.

#### Kidney

Kidney was dissected into three parts: renal cortex, renal medulla and renal pelvis. After washed in DMEM/F12 which added 10% FBS, these three parts of the kidney were minced respectively. 500μg/ml Liberase (TM) (Roche 5401119001) was firstly used to digesting the tissues into single cells at 37℃ for 40min, followed by 10 minutes of digestion in TryplE with shaking. In assistance with dissociation, pipette the cells during the incubation every 5 minutes. The dissociated cells were collected by centrifugation (800g, 5min), and further washed by DMEM/F12 added 10% FBS. Cells were resuspended and stained with 7-AAD before single-cell sorting. For METATAC, the kidney was dissected into three parts, renal cortex, renal medulla and renal pelvis. After dissection, large tissues were cut into small pieces by the blade in PBS, and transferred to a 40-um cell strainer, then were homogenized with the rubber tip of a syringe plunger (5ml) in 4ml PBS. The filtered cells were transferred to a 15ml tube and pelleted by centrifuge at 500g for 5min at 4c, then wash once with ice-cold PBS. All cells were cryopreserved in 90% fetal bovine serum and 10% DMSO.

#### Small intestine and Large intestine

After obtaining small intestine and large intestine from human embryo between 19w to 22w, we divided the small intestine into upper, middle and lower parts and the large intestine into transverse colon, ascendant colon and descendant colon parts. Then washed them by DMEM medium (plus 10% PBS) twice. Striated muscular layer and cut up, then added 500ul enzyme mix (2.5mg/ml Collagenase II Invitrogen, 2.5mg/ml Collagenase IV Invitrogen and 0.1 mg/ml DNase I dissolved in DMEM medium), 37°C and 1000rpm for 30-50min. Then added 500ml DMEM medium (plus 10% PBS) and used 40um Pre-Separation Filters filter the cell suspension. Tissues that were not fully digested were redigested with TrypLE. After centrifuging cell suspension at 800g, 5min, added 200ul PBS (plus 1% BSA) resuspended and added 5ul 7-AAD at room temperature for 10min. Then centrifuged cell suspension at 800g, 5min and used 500ul PBS (plus 1% BSA) resuspend.

#### Liver

We washed the liver sample twice with cold PBS to remove impurities and fat mass, then divide the liver into eight functionally independent segments (Segment I-VIII), each segment with its blood vessels and bile circulation. Next, each sample was fully minced with surgical scissors. We added the digestion buffer (2.5mg/ml II collagenase, 2.5mg/ml IV collagenase, 0.1 mg/ml DNase I in DMEM) and incubated the mixture at 37°C with shaking. We checked the proportion of single cells under the microscope every 10 minutes and the entire digestion process was up to 90 mins. We stopped the digestion procedure when the suspension contained 80-100% single cells. Cells were then filtered through a 40 µm strainer, pelleted (800g, 5 minutes), resuspended with Red Blood Cell Lysis Buffer, and kept at room temperature for 5 minutes to remove red blood cells. We then centrifuged (800g, 5min) and washed the pellet once with PBS. Finally, the cells were resuspended in FACS buffer, stained with 7-AAD and sorted by FACS.

#### Lung

Lung tissues were digested as two parts, lung center and lung peripheral. Both were firstly washed by DMEM containing 10% FBS, and then transferred into tubes containing 1mL of collagenase Ⅱ/collagenase Ⅳ (2.5 mg/ml). The tubes were incubated at 37℃ for ten minutes with shaking at 1000 rpm for digestion. Then the digested tissues were centrifuged to get cell pellets, which were resuspended in DMEM containing 10% FBS later. Samples were filtered, centrifuged, and dyed as described in bronchus sample collection. We collected 2000 cells for each part.

#### Testis and Ovary

Human gonad tissues include testis and ovary were obtained from human embryo from 19w to 22w. Washed the gonad tissues with DMEM medium (plus 10% PBS) twice and cut them up. Then added 600ul Accutase Cell Detachment Solution (Millipore #SCR005) at 37 °C, 1000rpm for 15min. then used 40um Pre-Separation Filters to filter the cell suspension. After centrifuging cell suspension at 800g, 5min, added 200ul PBS (plus 1% BSA) resuspended and added 10ul KIT FACS antibody at 4 °C for 30min. Then centrifuged cell suspension at 800g, 5min and used 200ul PBS (plus 1% BSA) resuspend. After that added 5ul 7-AAD at room temperature for 10min and centrifuged at 800g, 5min. Then used 500ul PBS (plus 1% BSA) resuspend.

#### Pancreas

Pancreas was processed to single-cell isolation immediately after the separation from the embryo. DMEM/F12 with 10% FBS was used to wash the pancreas for at least three times. The pancreas was sequentially minced using scissors and digested with dissociation buffer which containing collagenase Type II/IV (GIBCO, 17101015, 17104019) mixture and DNase I (Roche, 10104159001). After 30 minutes of incubation at 37℃, the digested cells were pelleted (800g, 5 minutes), washed in DMEM/F12 with 10% FBS once, passed through 40μm strainer, pelleted again. The cells were stained with 7-AAD for the assessment of cells’ viability before sorting.

#### Spleen

Spleen was cut into pieces and ground through a 40um strainer with syringe plunger in resuspension buffer (DMEM + 10% FBS). After centrifugation, cells were treated with ACK lysis buffer for 5min at 25℃ twice, centrifuged, and washed twice with resuspension buffer. Cells were stained with 1:40 7-AAD subsequently for FACS sorting.

#### Stomach

After removing the muscle layer with tweezers under the stereomicroscope, the stomach sample was divided into three parts, namely, fundus, body and antrum. Next, we stripped the fatty layer and blood vessels of the sample, washed with cold PBS 2-3 times to remove mucus, minced in a centrifuge tube and added with digestion buffer (2.5mg/ml type II/ IV collagenase, 0.1 mg/ml DNase I, in DMEM). After digestion 30 to 50mins at 37℃ with multiple pipetting to promote digestion procedure, cells were filtered through a 40 µm strainer, centrifuged at 800g for 5min, washed once with resuspension buffer (DMEM + 10% FBS) and pelleted again (800g, 5min). Finally, cells were resuspended with PBS containing 0.1% BSA and stained with 7-AAD.

#### Thymus

The thymus samples were crushed on a 100 µm strainer. Cells were centrifuged (500g, 5 minutes), digested with digestion buffer (2.5mg/ml II collagenase, 2.5mg/ml IV collagenase, 0.1 mg/ml DNase I, in DMEM) and incubated at 37°C for 30 minutes with agitation. The digestions quenched with resuspension buffer (DMEM + 10% FBS). Cells were pelleted (800g, 5 minutes), then resuspended in FACS buffer, and stained with 7-AAD immediately before sorting.

### Single-cell RNA-seq experiment

RNA-Seq was performed by the method of MALBAC-DT (Chapman et al., 2020). To improve throughput and reproducibility, an automated workflow was developed by using the Biomek FXP Workstation. A single-arm system with multichannel pipettor was used for RNA amplification and a dual-arm system with multichannel pipettor and Span-8 pipettors was used for sequencing library preparation. During RNA amplification, plates were kept on PCR-Cooler while transferring liquid and vortexed and briefly centrifuged after all transferring steps. If the plate will be stored at −80℃, a foil film was used for sealing; otherwise, an adhesive film was used. In this study, the 96 RT3-An primers with a distinct primer corresponding to each well were used to eliminate the possibility of cross-contamination between wells.

First, 96-well single cell capture plates containing cell lysis buffer were prepared. Cell lysis buffer of 2500 reactions consisting of 612.5uL H2O, 1000uL 5x SSIV buffer, 250uL 10% ICA-630, 2000uL 5M betaine, 125uL SUPERase•In RNase Inhibitor, 500uL 10mM dNTP mix and 12.5 uL 8×10^4^ diluted ERCC RNA Spike-In mix were prepared in a 5mL Eppendorf tube and distributed to three 8-strip tubes with 187uL in each well manually. Next, 45uL of the mix was transferred to each well of a 96-well master mix plate and then a transfer of 5uL 50uM barcoded RT-An primer from the primer storage plate to the master mix plate by the robot. After that, 2ul lysis buffer was distributed to each well of 24 capture plates from the master mix plate automatically and then stored at −80℃. Before cell sorting, capture plates were thawed at 4℃ and spun down for 15 seconds to collect the lysis buffer to the bottom of the well. After cell sorting, plates were spun down for another 15 seconds to ensure cell immersed into the lysis buffer and immediately stored at −80℃ until ready for amplification.

To perform reverse transcription, captured plates were incubated at 72°C for 3 minutes and hold at 4℃ to facilitate the open of RNA secondary structure and annealing of RT-An primer. RT mix of 2230 reactions consisting of 1807uL H2O, 892uL 5x SSIV buffer, 446uL 100mM DTT, 335uL SUPERase•In RNase Inhibitor, 536uL 100mM MgSO4 and 446uL SuperScript IV were prepared in a 5mL Eppendorf tube and distributed to three 8-strip tubes with 185uL in each well manually. Next, 45uL of RT mix was transferred to each well of a 96-well master mix plate by robot and 2ul RT mix were distributed to each well of 20 captured plates automatically. Incubate plates at 55°C for 10minutes to synthesize first strand cDNA.

After reverse transcription, excess RT primers were digested by exonuclease I, RT-Bn primers were added for an indication of digestion efficiency in this step. Exonuclease mix of 2230 reactions consisting of 2230uL H2O, 446uL ExoI buffer, 1338uL ExoI were prepared in a 5mL Eppendorf tube and distributed to three 8-strip tubes with 167uL in each well manually. Next, 41.4uL of the mix was transferred to each well of a 96-well master mix plate and then a transfer of 4.6uL 50uM barcoded RT-Bn primer from the primer storage plate to the master mix plate by the robot. After that, 2ul exonuclease mix was distributed to each well of 20 sample plates automatically and incubate plates at 37°C for 30 minutes to digest excess primers then at 80°C for 20 minutes to inactive exonuclease I.

For cDNA amplification, PCR master mix of 2000 reactions consisting of 38.48mL H2O,6mL ThermoPol buffer, 800uL 10mM dNTP mix, 320uL 100mM MgSO4, 200uL 200uM GAT-7N, 200uL 200uM GAT-COM and 2000uL Deep Vent (exo-)) were prepared in a 50mL tube and 250uL were added to each well of a 96-well master mix plate with an Eppendorf Multipette® E3 pipetter. Then, 24ul PCR master mix was distributed to each well of 20 sample plates from the master mix plate using the robot. PCR amplification conditions were as described in MALBAC-DT protocol but the cycles for exponential PCR were decreased from 18 to 15.

Finally, 2uL 10uM Tru2-G-RT primer was added to each well of the sample plates by robot and running an additional 5 cycles of PCR steps according to MALBAC-DT protocol.

Before sequencing library preparation, 5uL from each well of a sample plate was pooling to a 1.5mL tube automatically by Span-8 pipettors for one library preparation. After pooling, 50ul from the pooled samples were transferred to a 96-well plate and purified using 0.8x Ampure Beads. Next, the purified products were quantified using FilterMax F3 plate reader and 50ng DNA input was used for library preparation. Illumina sequencing adapters were added by tagmentation following manufacturer’s instructions of Vazyme TruePrep DNA Library Prep Kit. The PCR cycling conditions were as follows: 72℃ for 5 min; 98℃ for 30 sec; 12 cycles of 98℃ 10 sec,63℃ 30 sec, 72℃ 1 min; 72℃ for 5min. During PCR steps, Illumina Truseq read2 (Tru-R2) primers and Nextera 5XX primers were used to selectively amplify the 3’ ends of transcripts containing cell barcodes and UMIs. Paired-end sequencing was performed on an Illumina NovaSeq 6000 using 2 x150bp reads with a custom sequencing primer for read2. For a specific S4 run, 48 samples were sequenced with 12 samples multiplexed in each lane.

### Single-cell ATAC-seq experiment

#### Nuclei extraction

Quick thaw two tubes of each tissue cells at 37℃ water bath, then wash once with ice-cold PBS, count cell number, aliquot 50,000 to 1.5ml PCR tube (Eppendorf), centrifugation at 500xg for 5min at 4℃ with a swing bucket centrifuge. Nuclei were extracted with Omni-ATAC protocol (Corces et al., 2017), add 50ul ice-cold cell lysis buffer (10mM Tris, ph7.5, 10mM NaCl, 3mM MgCl_2_, 0.01% digitonin, 0.1 IGEPAL CA630, 0.1% Tween 20), pipette to mix thoroughly, put on ice for 3min, then add 100ul ice-cold wash buffer (10mM Tris, ph7.5, 10mM NaCl, 3mM MgCl_2_, 0.1% Tween 20), pelleted by centrifuge at 500xg for 10min at 4℃, wash once with 100ul ice-cold wash buffer, pelleted nuclei.

#### Assemble META transposome

We use META transposome (Tan et al., 2018) in the transposition step, to avoid half loss as compared to the Nextera transposome. One strand of the transposon was 5′-/Phos/-CTGTCTCTTATACACATCT-3′, while the other strand was in the form of 5′-[META tag]-AGATGTGTATAAGAGACAG-3′. Each of the oligos (Invitrogen, purification: PAGE) was dissolved in 0.1 X TE to a final concentration of 100 uM. For each of the *n* = 16 META tags, two strands were annealed at a final concentration of 5 uM each. The 16 annealed transposons were then pooled with equal volumes. The transposase was purified after expression from the pTXB1-Tn5 plasmid (Addgene). Transposome was assembled at a final concentration of 1.25 uM dimer (2.5 uM monomer).

In this work, we use META with n=16 tags:

CGAGCGCATTAA

AGCCCGGTTATA

TCGGCACCAATA

GCCTGTGGATTA

GCGACCCTTTTA

GCATGCGGTAAT

GCGTTGCCATAT

GGCCGCATTTAT

ACCGCCTCTATT

CCGTGCCAAAAT

TCTCCGGGAATT

CCGCGCTTATTT

CTGAGCTCGTTTT

#### Transposition

Resuspend pellet in 25ul transposition mix (12.5ul 2x TD buffer from Nextera kit, 10ul PBS ph7.5, 0.25ul 1% Digitonin, 0.25ul 10% Tween, 2ul 1.25uM META transposome), pipette to mix thoroughly, then incubate in a thermomixer at 1000rpm for 30min at 37℃. After transposition, add 25ul 2 x STOP buffer (40mM EDTA, 10mM Tris pH 8.5, 1mM spermidine), incubate on ice for 15min to stop transposition.

#### FACS single nuclei and amplification

For FACS, resuspend transposed cells in 1.5ml 0.5% BSA in PBS, then sorted single cells into 96-well plates containing 1ul lysis buffer (10mM Tris pH 8.0, 20mM NaCl, 1mM EDTA, 0.1% SDS, 500nM Carrier ssDNA, 60ug/ml QIAGEN protease) with a BD flow cytometer (BD, AriaII). Events were first gated on FSC and SSC as “cells”, and then on FSC and trigger pulse width as “singlets”. The sorting mode was “1.0 drop single”. After sorting, plates were sealed with an aluminum sealing film (PCR-AS-600, Axygen), centrifugation at 2800xg with swing bucket centrifuge for 1min at 4℃ to ensure nuclei in lysis buffer, then store at −80℃ until ready for PCR amplification.

We thawed plates, change with an adhesive sealing film (MSB1001, bio-rad), then incubate at 65℃ for 15min to release Tn5 from DNA on a thermocycler, then add 1ul 3% Triton X-100 to quench SDS. For amplification, first add 4ul preamp mix (3ul 2x high fidelity Q5 Master mix, 0.192ul 50uM META16 primer mix, 0.05ul 100mM MgCl_2_, 0.758ul H_2_O) to each well, cycling conditions were as follows

• 72℃, 5min,

• 98℃, 30s

• 16 cycles:

98℃, 10s

62℃, 30s

72℃, 1min

• 72℃, 5min

• hold at 4℃

After preamplification, add 0.225ul 50uM indexed META16-ADP1 primer to each column, and 0.225ul 50uM META16-ADP2 primer to each row to incorporate well-specific cell barcodes, cycling conditions were as follows

• 98℃, 30s

• 5 cycles:

98℃, 10s

62℃, 30s

72℃, 1min

• 72℃, 5min

• hold at 4℃

After amplification, pool a whole plate, purify with DNA Clean & Concentrator-5 column (ZYMO).

META16 primer mix sequence in the form of 5’-[META tag]-AGATGTGTATAAG META16-ADP1 primer design in the form of 5’-CTTTCCCTACACGACGCTCTT CCGATCT-[Cell Barcode]-[META Tag]-AGATGTGTATAAG. META16-ADP2 primers design in the form of 5’-GAGTTCAGACGTGTGCTCTTCCGATCT-[Cell Barcode]-[META Tag]-AGATGTGTATAAG.

ADP1 cell barcodes as follows

GATATG, ATACG, CCGTCTG, TGCG, GAACTCG, ATGTAG, CCCG, TGTAG, GAGTAAG, ATCG, CCTAG, TGACCG

ADP2 cell barcodes as follows

ACTCTA, AGAGCAT, GGTATG, TCGATGC, CTACTAG, TATGCA, CACACGA, GTCGAT

All liquid transfer steps were handled by a liquid handler platform Echo525.

Detailed calibration for each transfer steps is as follows:

Cell lysis buffer: 384PP_AQ_BP

3% Triton X-100: 384PP_AQ_SPHigh

PCR master mix: 384PP_AQ_BP

Primer mix: 384PP_AQ_BP

#### Library preparation and sequencing

Each plate takes 120ng (9ul template) for library preparation, the library was performed by addition of 21ul PCR mix (15ul 2x Q5 Master mix, 3ul NEBNext index primer i5, and 3ul NEBNext index primer i7, 0.05ul 100mM MgCl2), here we use a unique dual index combination for each plate to reduce index hopping. Then incubate as 98℃, 30s, 2 cycles [98℃, 10s, 68℃, 30s, 72℃, 1min], note only 2 cycles to avoid cell barcode switching in case of any remaining cell barcode primers. Finally, the library concentration was determined by Kapa qPCR master mix. For sequencing, the equimolar libraries from each 96-well plate were pooled and sequenced on two runs of a NovaSeq 6000 (Illumina).

### Single-cell RNA-seq data analysis

#### Data pre-processing

First, the raw data for each 96-well plate was demultiplexed according to the cell barcodes in the R2 reads, where no mismatch was allowed. Then, the demultiplexed R2 reads which contained less than 3 bases inconsistent to designed UMI patterns and contained at least 4 T in the 5bp downstream regions of UMIs were recognized, and the corresponding R1 reads were extracted. For the remained R1 reads, the polyA sequences were trimmed, followed by filtering for high-quality reads with the following criteria: 1) at least 40bp; 2) more than half of the bases showing the sequence quality scores greater than 38. 3) less than 10% of bases showing N.

#### Reads mapping and gene expression calculation

The processed R1 reads were mapped to the human genome (GRCh38.p10) using HISAT2 (Kim et al., 2015) with the option of “--new-summary”, where the genome and gene annotation files (primary assembly) were download from Gencode (Frankish et al., 2019). The reads mapped to multiple genomic positions were removed according to the “NH:i” tags in the BAM files using Samtools (Li et al., 2009). The remained reads were assigned to genes using htseq-count in HTSeq (Anders et al., 2015) with the default options. Then for each gene, the reads with similar UMIs (no more than 2 hamming distance) were collapsed to remove redundant reads. Finally, the UMI count for each gene was calculated to generate the gene expression matrix.

#### Cell and gene filtering

Several criteria were used for cell filtering in each organ: 1) the ratio of primer A in all primers (A and B) >= 0.9; 2) clean reads number >= 0.4 million; 3) reads mapping ratio >= 0.6; 4) detected gene number > 1000; 5) UMI number > 3000. 6) mitochondrial gene UMI ratio < 0.15; 7) ERCC ratio < 0.25. To filter out doublets, the cells showing extremely high gene number and UMI number were removed. In addition, a generalized additive model (GAM) was fit for UMI number (y) against gene number (x) using gam in mgcv. The cells showing the observed UMI number great than 2-fold of the expected UMI number were removed. After cell filtering, the genes were filtered using two strategies: 1) in each organ, the genes expressed in less than 10 cells were removed, which produced the files used for analysis in each organ. 2) the gene expression matrices in each organ were merged and then the genes expressed in less than 10 cells were removed, which produced the file (47,468 genes by 31,208 cells) used for analysis across 17 organs.

#### Cell type identification

For each organ, the gene expression matrix after filtering was used for cell clustering using Seurat (Butler et al., 2018). Specifically, the highly variable genes were identified using FindVariableGenes with the options of “mean.function = ExpMean, dispersion.function = LogVMR, x.low.cutoff = 0.25, x.high.cutoff = 5, y.cutoff = 0.5”, followed by PCA dimension reduction. The significant dimensions with the p-value less than 0.001 were used for cell clustering. The resolution was optimized to produce reliable clusters according to the *t*-SNE plot. To avoid over-clustering, the similar cell clusters were merged using ValidateClusters with the options of “top.genes = 30, min.connectivity = 0.01, acc.cutoff = 0.85”. The differentially expressed genes (signature genes) in each cell cluster were identified using FindMarkers, and only the signature genes with power >= 0.4 and fold change >= 2 were selected. Based on signature genes, cell identities were assigned to each cell cluster. Gene ontology enrichment analysis was performed using runTest (Fisher’s exact test) in topGO (Alexa and Rahnenfuhrer, 2018), and only the terms with FDR < 0.05 were selected.

#### The comparison between fetal and adult cells

The single-cell RNA-seq data for the 6 organs (kidney, large intestine, liver, lung, spleen and testis) in the human adult was downloaded from Single Cell Portal (https://singlecell.broadinstitute.org/single_cell) and literature (Guo et al., 2018; Kinchen et al., 2018; MacParland et al., 2018; Madissoon et al., 2019; Stewart et al., 2019). The cells with at least 500 detected genes were used for analysis. For each organ, the adult cells were mapped into the fetal cells using Cell Blast (Cao et al., 2019b) with the cutoff of 0.2 for cell type identification. To remove the batch effect, the expression matrix of fetal and adult cells was corrected for each organ using Seurat (Stuart et al., 2019). To compare the heterogeneity between fetal and adult cells, pairwise Spearman correlation was calculated between the cells within each organ and each cell group (epithelial cells, endothelial cells, fibroblasts and immune cells) for fetal and adult data, respectively. To compare the distance between fetal and adult cells across cell groups, pairwise Spearman correlation was calculated between the fetal cells and the adult cells within each organ and each cell group.

#### Cross-organ analysis

To perform hierarchy clustering across organs, the gene expression matrix containing all the 17 organs was normalized into count per million (CPM). The cell types which belonged to the same identity were grouped. For example, all the epithelial cells in the kidney were grouped into kidney epithelial cells. For each cell group in each organ, the average CPM was calculated. Then, the pairwise Pearson correlation was calculated between cell types. Hierarchy clustering (average linkage) was performed based on the distances (1 - correlation).

To identify the putative TFs playing roles in cell type commitment, the CPM for TFs were extracted from the gene expression matrix containing all the 17 organs. Then random forest classification was performed using randomForest in randomForest with the option of “importance = T”. The TFs were decreasingly ordered by the mean decrease in Gini index.

#### Correlated gene module (CGM) detection and annotation

To estimate the required cell number for CGM analysis, the most abundant cell type (Fibro-ADAM28) was randomly subsampled into 100, 300, 500, …, 1300, 1500, 1700 cells, respectively. For each group of sampled cells, the gene expression was normalized into count per million (CPM) and pairwise gene correlation (Spearman) was calculated to generate the background distribution of gene correlation. The 95% quantile of correlation showed robust for the groups with at least 500 cells, thus 500 was used as the required cell number.

For each of the 10 cell types passing this requirement, 500 cells were randomly subsampled for gene correlation calculation as mentioned above. Hierarchy clustering (average linkage) was performed based on the distances (1 - correlation) and CGMs were detected using cutreeDynamic in dynamicTreeCut (Langfelder et al., 2008) with the options of “cutHeight = 0.99, minClusterSize = 10, method = “tree”, deepSplit = F”, and only the CGMs with the average correlation >= the 95% quantile of background correlation (0.088) were selected. Then, the CGMs in different cell types were merged. To remove redundancy, the pair of similar CGMs (the ratio of common genes number to union genes number > 0.6) was replaced by their common genes.

To calculate the gene type composition for each CGM, the gene type was extracted from the gene annotation file mentioned in reads mapping. The human TF list was downloaded from Animal TFDB v3.0 (Hu et al., 2019).

To perform enrichment analysis for the genes in each of the 227 CGMs with high speed, three types of datasets were built:

1. The dataset for TF enrichment. The binding motifs were downloaded from JASPAR (CORE) and TRANSFAC^®^, respectively. The human motifs were extracted and only the ones in the TF list mentioned above were selected. For the TFs with more than one motif, only the non-variants or the recommended one was chosen according to the motif annotation information. For the TFs existing in both JASPAR and TRANSFAC^®^, only the one in the former was chosen. Then, the gene promoter (upstream −2kb of TSS ∼ downstream of TSS of genes) sequences were extracted using BEDTools (Quinlan and Hall, 2010), followed by the Markov model estimation using fasta-get-markov in MEME (Bailey et al., 2009). After that, TF binding sites were identified using FIMO (Grant et al., 2011) in MEME with the options of “--parse-genomic-coord -- thresh 1e-5 --max-stored-scores 500000”. The genes whose promoter regions containing TFBSs were assigned to the target genes of the corresponding TFs.
2. The dataset for gene ontology (GO) enrichment The gene ontology annotation file in the human (goa_human.gaf.gz with the time stamp of 2019-10-09) was downloaded from The Gene Ontology Consortium (The Gene Ontology Consortium, 2019). The terms in the biological process aspect were extracted.
3. The dataset for KEGG pathway enrichment The KEGG pathway files in the human (with the time stamp of 2019-12-18) were downloaded from the KEGG database (Kanehisa et al., 2017). These three datasets were then processed and used for gene enrichment analysis using enricher in clusterProfiler (Yu et al., 2012) with the options of “pAdjustMethod = “BH”, minGSSize = 1, maxGSSize = Inf, pvalueCutoff = 0.05”, and the results with the adjusted p-value less than 0.05 were selected.

In addition, the protein-protein interaction (PPI) enrichment was performed using STRINGdb (Szklarczyk et al., 2015), where the PPI data was imported using STRINGdb$new with the options of “version = “10”, species = 9606, score_threshold = 400” and enrichment test was performed using STRINGdb$get_ppi_enrichment. The results with the p-value less than 0.05 were selected.

### Single-cell ATAC-seq data analysis

#### Data pre-processing and quality control

First, the raw data for each 96-well plate was demultiplexed according to the cell barcodes in the R1 and R2 reads, where at most 1 mismatch was allowed. Then, reads were trimmed using cutadapt (Martin, 2011) with the option “-e 0.22” to remove cell barcodes, META tags and transposon sequences at both 5’ and 3’ ends. The trimmed paired-end reads were aligned to the human genome (GRCh38.p10) using Bowtie2 (Langmead and Salzberg, 2012) with “-X 2000 --local --mm --no-discordant --no-mixed” parameters. PCR duplicates and contaminated reads were subsequently removed for reads that aligned to the autosome and X chromosome with mapping quality >= 20 using a custom python script based on the coordinates and META tags. The cell filtering was based on the following criteria: 1) number of reads >= 100k; 2) number of unique fragments >= 10k and <= 600k; 3) ratio of reads aligned to the nuclear genome >= 0.85; 4) ratio of reads aligned to the mitochondria <= 0.1.

#### Peak calling and feature matrix

The unique fragments of all cells passing quality control from 14 organs were merged together. Accessible peaks were called on the merged files using macs2 callpeak command (Zhang et al., 2008) with “-f BEDPE --nomodel --nolambda --SPMR --keep-dup all” as options. Peaks overlapped with the ENCODE blacklist were removed. Peaks with size longer than 2kb were broke into ∼1kb windows. 333,614 accessible peaks were identified. For each cell, the accessibility of each peak was quantified by the count of Tn5 insertion which occurred within this peak to construct the feature matrix.

#### TF binding site identification

We downloaded human TF motif position weight matrices from JASPAR core database (2018) and TRANSFAC^®^, respectively. The peak sequences were extracted using BEDTools (Quinlan and Hall, 2010), and the motif binding sites on each accessible peaks were identified using FIMO with parameters “--thresh 0.0001 --max-stored-scores 500000 --max-strand”.

#### TF motif accessibility score

We used chromVAR (Schep et al., 2017) to calculate the global TF motif accessibility deviations. The peak-to-cell feature matrix and genomic coordinates were input to “SummarizedExperiment” function to construct the object. “addGCBias” function was applied to compute the GC content for peaks. The the JASPAR 2018 core database was got using “getMatrixSet” function with “‘species’ = 9606, ‘all_versions’ = T” as options, and mapped to hg38 genome by “matchMotifs” function with option “genome = BSgenome.Hsapiens.UCSC.hg38”. The TF motif accessibility deviations were calculated using “computeDeviations” with default parameters based on the motif annotation and the “SummarizedExperiment” object.

#### Co-accessibility and gene activity scores

We used R package Cicero (Pliner et al., 2018) to estimate gene activities from METATAC data. In brief, the sparse binary cell-by-peak matrix of each organ was used to create a cell_data_set. After “detect_genes” and “estimate_size_factors”, we performed Latent Semantic Indexing (LSI) and reduced dimensions by UMAP. The UMAP coordinates were input to “make_cicero_cds” function, and then “run_cicero” to get co-accessibility scores between peaks with default parameter “k = 50”.

After got the co-accessibility scores between peaks, we used “annotate_cds_by_site” function to annotate all peaks located in the gene body +2kb upstream of TSS, and then used “build_gene_activity_matrix” with default parameter “dist_thresh = 250000, coaccess_cutoff = 0.25” to calculate gene activity scores for each gene in each cell.

#### Cell type identification using Seurat’s canonical correlation analysis

We applied Seurat’s scATAC-seq + scRNA-seq integration pipeline to match the cellular states across two modalities, and transfer cell labels from transcriptomic data to chromatin accessibility data (Stuart et al., 2019). For each organ, scATAC-seq gene activity score matrix and annotated gene expression matrix were normalized and scaled, and then input to “FindTransferAnchors” function with “reduction = ‘cca’” as the option to generate anchor set between two datasets. This anchor set was input to “TransferData” function, weighted by the LSI reduced scATAC-seq peak-to-cell matrix, to predict cell identity of scATAC-seq cells.

#### Cross-organ analysis

The peak-to-gene count matrix and the gene activity score matrix containing 14 organs were log-normalized and scaled using Seurat “NormalizeData” and “ScaleData” functions. The cell types which belonged to the same identity were grouped, as mentioned in “Single-cell RNA-seq data analysis” part. For each cell group in each organ, the average scaled accessibility of each peak and average scaled gene activity score of each gene were calculated. We also calculated the *Z* scores for the TF motif accessibility deviation matrix of 14 organs, and then took the average for each cell group in each organ.

To compare the gene expressions and gene activity scores, we computed the Spearman correlation between average scaled expressions and average scaled gene activity scores for common cell types of both data.

To compare the TF expressions and TF motif accessibility scores, we computed the Spearman correlation between average scaled TF expression and average scaled TF motif accessibility scores for common cell types of both data.

To perform hierarchy clustering across organs, the pairwise Pearson of average scaled peak accessibilities correlation was calculated between cell types. Hierarchy clustering (complete linkage) was performed based on the distances (1 - correlation).

#### Regulatory regions for each gene

We re-calculated the co-accessibility and gene activity scores using Cicero for each cell type with more than 100 cells, to get cell-type-specific co-accessibility scores. For each gene in each cell type, the regulatory regions included the peaks located in the gene body +2kb upstream of TSS, and co-accessible peaks with co-accessibility score >= 0.25 and within 25kb.

#### Comparison between co-expression and co-accessibility

The co-expression index of two genes is defined as the Spearman correlation coefficient of their normalized gene expressions. The co-accessibility index of two genes is defined as the Jaccard index of their binary gene activity scores, which is the number of cells both gene activity score > 0, divided by the number of cells at least one of the two genes with activity score > 0.

The co-expression/co-accessibility index of a CGM is defined as the average co-expression/co-accessibility indexes of all gene pairs within the corresponding CGM, respectively.

#### Infer functional TFs for CGMs

For each CGM, we combined all regulatory peaks of its genes, and computed motif enrichment false discovery rate (FDR) using the hypergeometric test for JASPAR and TRANSFAC^®^ motifs. For TF with multiple motifs, we only kept the one with minimum FDR.

On the other hand, for each CGM and each TF, we calculated the average co-expression index of TF and each gene.

### Data and Code Availability

DATA RESOURCES: The accession number of the raw data files for the RNA-seq and ATAC-seq experiment reported in this paper is X. To take full advantage of the resource in the GeACT project, we made our data available for further exploration via an interactive website at http://geact.gao-lab.org.

SOFTWARE: All software is freely or commercially available and is listed in the STAR Methods description and Key Resources Table. The code is accessible at https://github.com/gao-lab/GeACT.

## Supplemental Information

Supplemental Information includes seven figures and four tables and can be found with this article online.

## Supplementary Figure Legends

**Figure S1.**
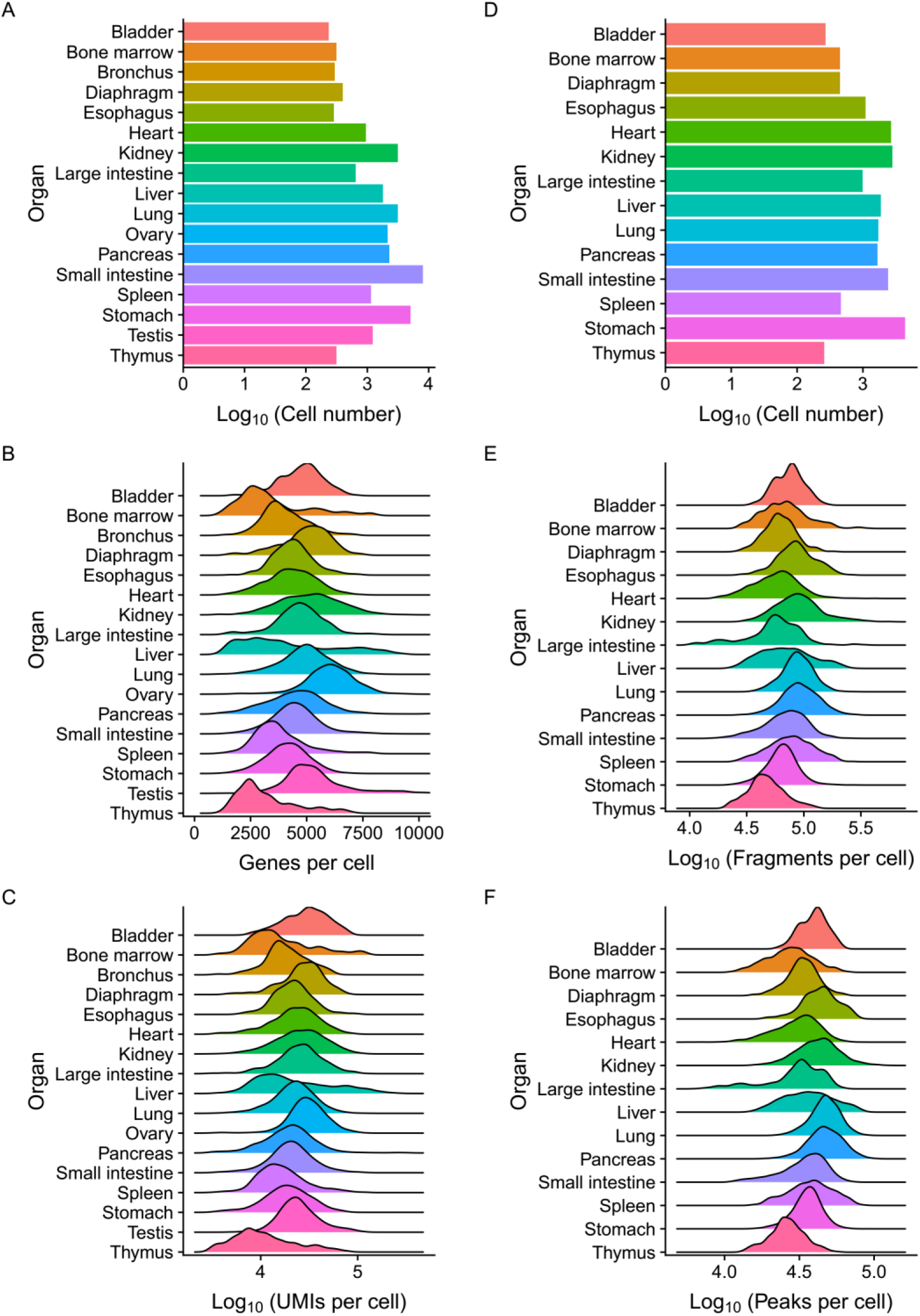
The overview of GeACT project, Related to. Figure 1. (A) The bar plot shows the number of cells after quality control for each organ in the single-cell RNA-seq data. (B and C) The distribution of detected gene number (B) and UMI number (C) in the single-cell RNA-seq data. (D) The bar plot shows the number of cells after quality control for each organ in the single-cell ATAC-seq data. (E and F) The distribution of detected fragment number (E) and peak number (F) in the single-cell ATAC-seq data.

**Figure S2.**
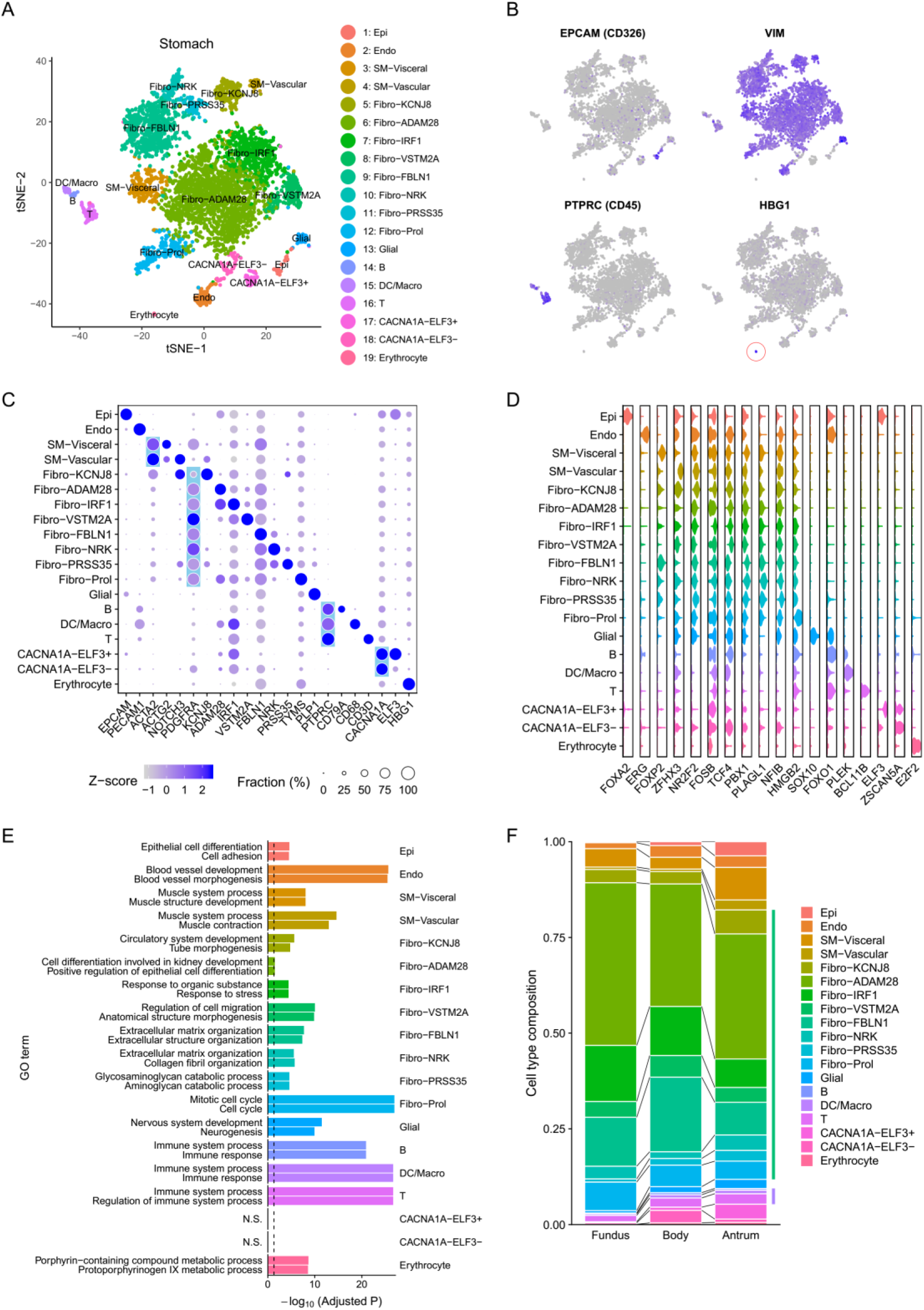
The single-cell transcriptome landscape in the stomach, Related to. Figure 2. (A) The *t*-SNE plot shows the cells colored by cell types. Epi: epithelial cells. Endo: endothelial cells. SM: smooth muscle cells. Fibro: fibroblasts. DC/Macro: dendritic cells or macrophages. (B) The relative expression level of marker genes for primary cell type groups. (C) The relative expression of signature genes in each cell type. (D) The relative expression of TFs in each type. Only the top 1∼2 significant TFs are shown. (E) The bar plot shows the enriched terms of GO biological process. Only the top 10 significant terms are shown. (F) The bar plot shows the composition of each cell type.

**Figure S3.**
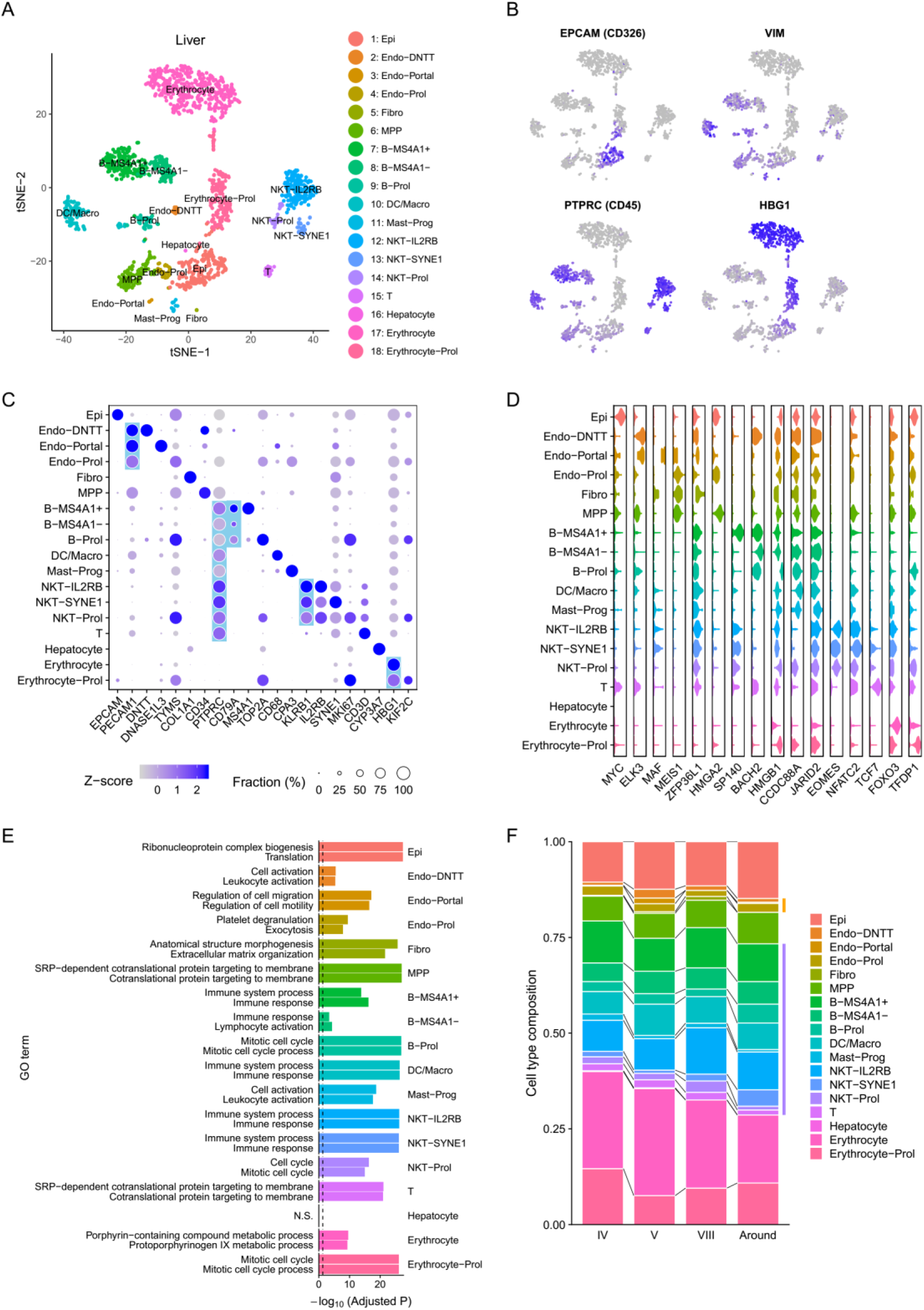
The single-cell transcriptome landscape in the liver, Related to Figure **2.** (A) The *t*-SNE plot shows the cells colored by cell types. Epi: epithelial cells. Endo: endothelial cells. Fibro: fibroblasts. MPP: multipotent progenitors. DC/Macro: dendritic cells or macrophages. “Prog” and “Prol” mean progenitor cells and proliferative cells, respectively. (B) The relative expression level of marker genes for primary cell type groups. (C) The relative expression of signature genes in each cell type. (D) The relative expression of TFs in each type. Only the top 1∼2 significant TFs are shown. (E) The bar plot shows the enriched terms of GO biological process. Only the top 10 significant terms are shown. (F) The bar plot shows the composition of each cell type. IV, V and VIII mean the corresponding segments. Around means the segments VII, VI, II and III.

**Figure S4.**
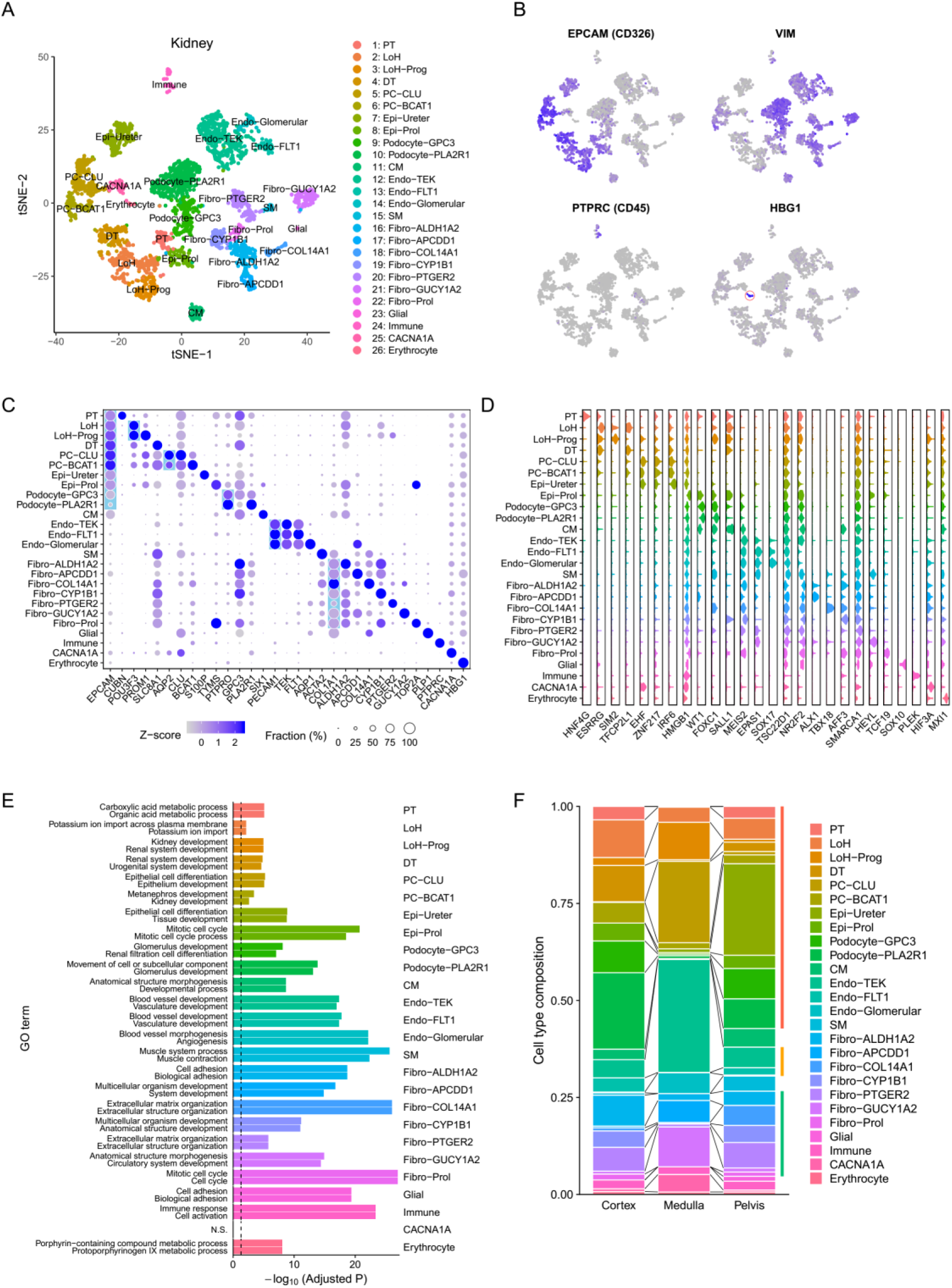
The single-cell transcriptome landscape in the kidney, Related to Figure **2.** (A) The *t*-SNE plot shows the cells colored by cell types. PT: proximal tubule. LoH: loop of Henle. DT: distal tubule. PC: principal cells. Epi: epithelial cells. CM: cap mesenchyme. Endo: endothelial cells. SM: smooth muscle cells. Fibro: fibroblasts. “Prog” and “Prol” mean progenitor cells and proliferative cells, respectively. (B) The relative expression level of marker genes for primary cell type groups. (C) The relative expression of signature genes in each cell type. (D) The relative expression of TFs in each type. Only the top 1∼2 significant TFs are shown. (E) The bar plot shows the enriched terms of GO biological process. Only the top 10 significant terms are shown. (F) The bar plot shows the composition of each cell type.

**Figure S5.**
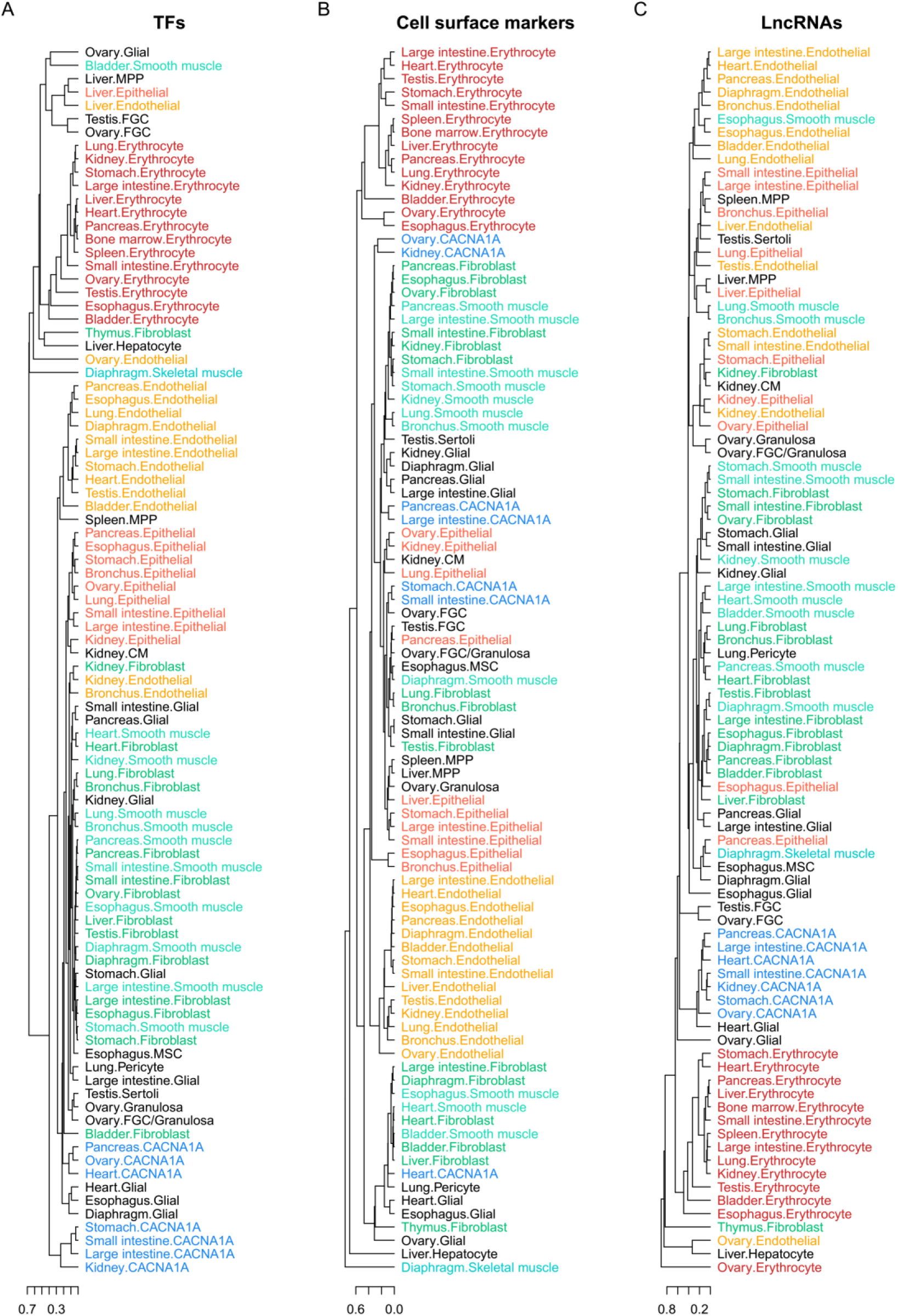
The architecture of expression profiles across cell types, Related to. Figure 3. The dendrograms show the clustering of gene expression for non-immune cell types based on TFs (A), cell surface markers (B) and lncRNAs (C), colored by cell types.

**Figure S6.**
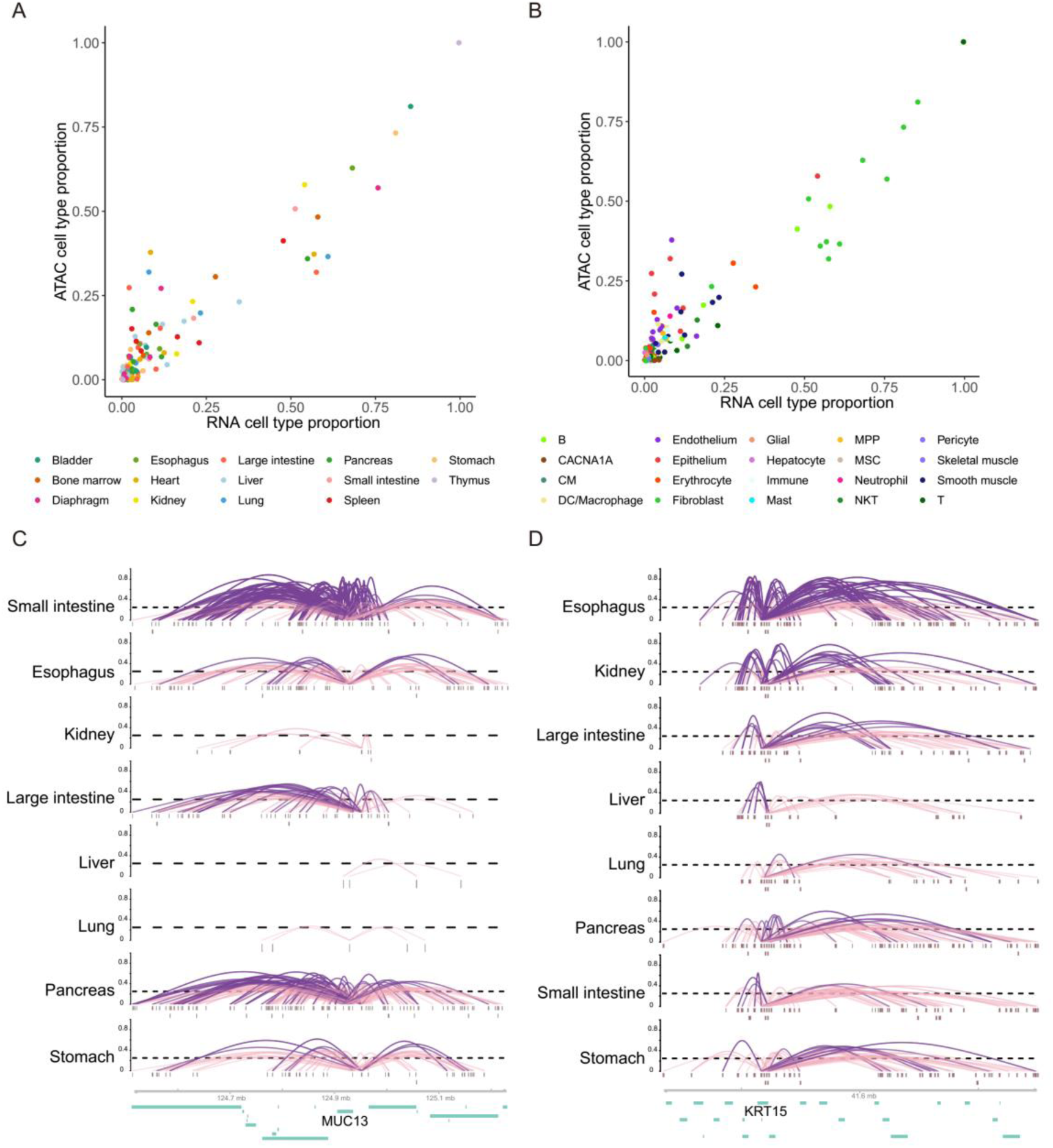
The architecture of single-cell chromatin accessibility profiles across organs and cell types, Related to. Figures 4 and 5. (A) Comparison of cell type proportion in each organ between METATAC data and MALBAC-DT, colored by organs. (B) The same as (A), colored by cell types. (C and D) Cicero peak-to-gene connections for the marker gene locus in different organs. Only connections with co-accessibility score >= 0.25 are shown. Connections with co-accessibility score >= 0.4 are colored by purple. (C) is for MUC13, which is a marker gene of epithelial cells in the small intestine. (D) is for KRT15, which is a signature gene of epithelial cells in the esophagus.

**Figure S7.**
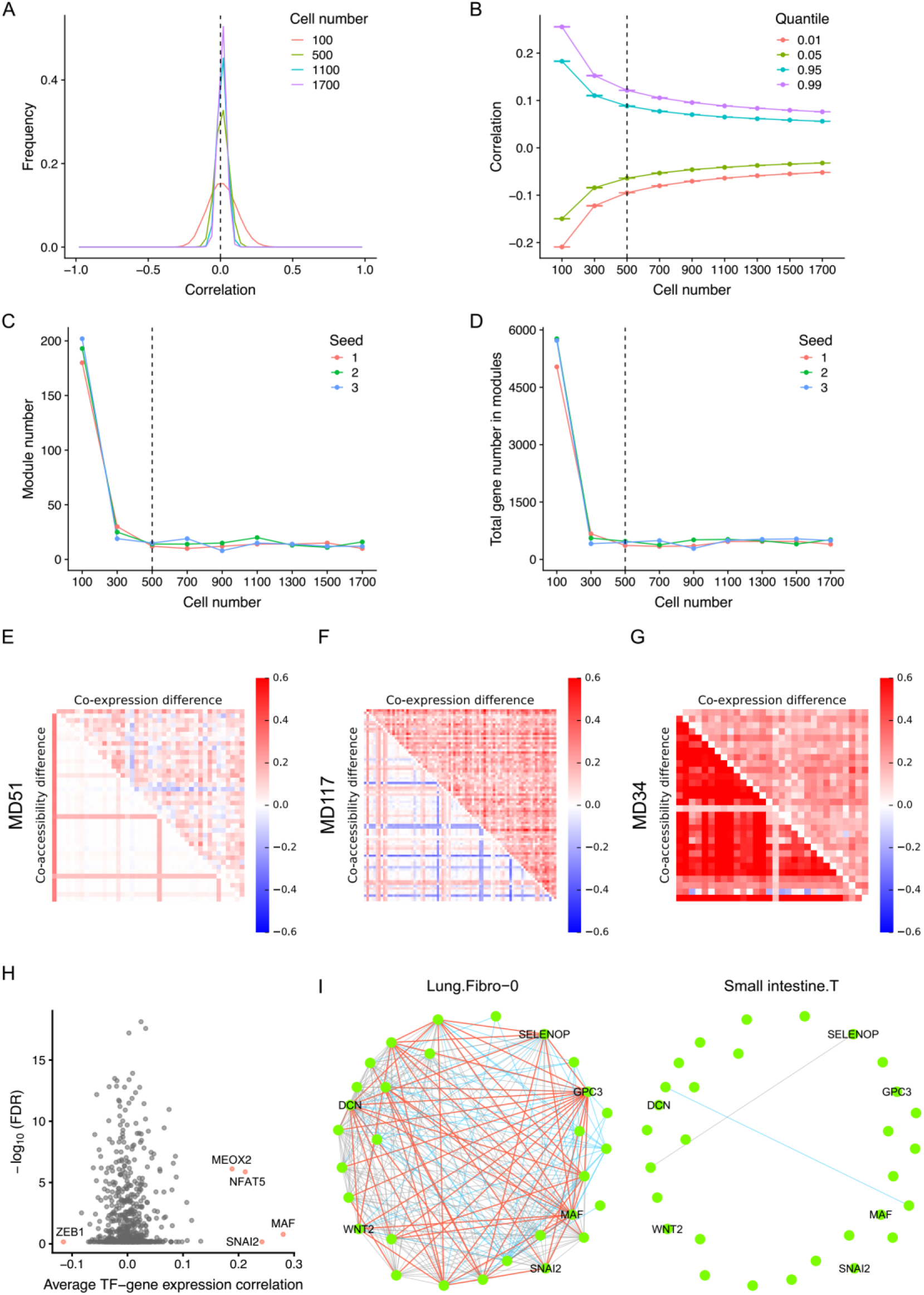
The co-expressed gene modules (CGMs) map across cell types, Related to. Figures 6 and 7. (A) The distribution of Spearman correlation for all gene pairs using 100, 500, 1100 and 1700 cells, respectively. (B) The quantile of correlation using 100∼1700 cells, respectively. (C) The number of detected gene modules using 100∼1700 cells, respectively. (D) The number of genes in gene modules using 100∼1700 cells, respectively. (E-G) The heatmap of co-expression differences and the co-accessibility differences of CGM genes between two cell types. The upper triangle is the expression Spearman correlation in cell type 1 minus that in cell type 2, and the lower triangle is the binary gene activity Jaccard index in cell type 1 minus that in cell type 2. E: MD51, pancreas Fibro-PAMR1+SOX6+ cells compared with stomach Fibro-FBLN1 cells. F: MD117, small intestine SM-Visceral cells compared with small intestine Fibro-COL6A5 cells. G: MD34, lung Fibro-0 cells compared with small intestine T cells.

## Supplementary Table Legends

Table S1. The summary of the samples and oligonucleotides used for sequencing, Related to Figure 1.

Table S2. The metatable and signature genes for the single-cell RNA-seq data, Related to Figure 2.

Table S3. The metatable for the single-cell ATAC-seq data, Related to Figure 4.

Table S4. The metatable for correlated gene modules (CGMs), Related to Figure 6.

